# An Integrated Mechanistic Model of Pan-Cancer Driver Pathways Predicts Stochastic Proliferation and Death

**DOI:** 10.1101/128801

**Authors:** Mehdi Bouhaddou, Anne Marie Barrette, Rick J. Koch, Matthew S. DiStefano, Eric A. Riesel, Alan D. Stern, Luis C. Santos, Annie Tan, Alex Mertz, Marc R. Birtwistle

## Abstract

Most cancer cells harbor multiple drivers whose epistasis and interactions with expression context clouds drug sensitivity prediction. We constructed a mechanistic computational model that is context-tailored by omics data to capture regulation of stochastic proliferation and death by pan-cancer driver pathways. Simulations and experiments explore how the coordinated dynamics of RAF/MEK/ERK and PI-3K/AKT kinase activities in response to synergistic mitogen or drug combinations control cell fate in a specific cellular context. In this context, synergistic ERK and AKT inhibitor-induced death is likely mediated by BIM rather than BAD. AKT dynamics explain S-phase entry synergy between EGF and insulin, but stochastic ERK dynamics seem to drive cell-to-cell proliferation variability, which in simulations are predictable from pre-stimulus fluctuations in C-Raf/B-Raf levels. Simulations predict MEK alteration negligibly influences transformation, consistent with clinical data. Our model mechanistically interprets context-specific landscapes between driver pathways and cell fates, moving towards more rational cancer combination therapy.

## INTRODUCTION

Oncogene-targeted small molecule kinase inhibitors, like imatinib for BCR-ABL (Druker et al., 2001), have transformed chemotherapy. However, such precision medicine approaches are not always efficacious. In some cases, mutation-matched patients do not respond to the drug (Cobleigh et al., 1999), or alternatively, resistance stochastically develops (Vogel et al., 2002). Monotherapy can even activate the target pathway, depending on cellular context (Poulikakos et al., 2010). Combination therapy is a logical and clinically-proven path forward (Flaherty et al., 2012), but rationalizing even just the choice of combinations from among the at least 28 FDA-approved (Wu et al., 2015) targeted small molecule kinase inhibitors, notwithstanding important questions related to the dozens of traditional chemotherapeutics, monoclonal antibodies, dose, timing and sequence, remains challenging for basic and clinical research. Approaches are needed that consider quantitative, dynamic, and stochastic properties of cancer cells given a context.

Computational modeling can help fulfill this need, since simulations often are much quicker and less expensive than the explosion of experimental conditions one would need to assay the drug combination space. Some of the first approaches define transcriptomic signatures that define tumor subtypes and suggest drug vulnerabilities (Sørlie et al., 2001). Big data and bioinformatic statistical approaches are at the forefront of predicting drug sensitivity, with penalized regression and other machine learning methods linking mutation or expression biomarkers to drug sensitivity (Barretina et al., 2012; Garnett et al., 2012).

Such statistical modeling approaches are often limited to extrapolating predictions into untrained regimes, which is important for drug combinations. They also usually cannot account for dose and dynamics, central to combinations. Alternatively, mechanistic computational models based on physicochemical representations of cell signaling have an inherent ability to account for dose and dynamics. They also have greater potential for extrapolation, since they represent physical processes. Multiple mechanistic models exist for almost every major pan-cancer driver pathway identified by The Cancer Genome Atlas (TCGA): receptor tyrosine kinases (RTKs); Ras/ERK; PI3K/AKT; Rb/CDK; p53/MDM2 (Ciriello et al., 2013). An integrative model that accounts a multi-driver context has not yet been built, but will likely be needed to address drug combination prediction problems with such modeling approaches.

Here, we constructed the first mechanistic mathematical model integrating these commonly mutated signaling pathways. We use the MCF10A cell line – a non-transformed mammalian cell line with predictable phenotypic behaviors. We tailor the model to genomic, transcriptomic, and proteomic data from MCF10A cells and train the model using a wealth of literature resources as well as our own microwestern blot and flow cytometry experiments to refine biochemical parameters and phenotypic predictions. We use the model to explore stochastic proliferation and death responses to pro- and anticancer perturbations. This mechanistic, biological context-tailored mathematical model depicting several major cancer signaling pathways allows us to probe the mechanisms that underlie how noisy signaling dynamics drive stochastic proliferation and death fates in response to various perturbations, and gain insight into their biochemical mechanisms.

## RESULTS

### Rationale and Workflow

An overarching goal of this work is a foundation that integrates canonical biochemical knowledge of cancer driver pathways with cell-specific context to make quantitative predictions about how drugs, microenvironment signals, or their varied combinations influence cell proliferation and death (Fig. 1A, S1; Table S1-S2). Three main considerations guided us. First is scope. TCGA identified pan-cancer driver pathways (Ciriello et al., 2013): multiple RTKs; proliferation and growth (Ras/Raf/MEK/ERK; PI-3K/PTEN/AKT/mTOR); cell cycle; DNA damage. These interface with expression and apoptosis (Fig. 1A). Second is inter- and intra-tumoral heterogeneity. Experimental accessibility defines each, which predominantly consists of copy number variation, mRNA expression and mutations, which the foundation integrates. Intra-tumoral heterogeneity is genetic (Landau et al., 2013) and non-genetic (Spencer et al., 2009), so we require accounting for such noise. Third is formalism. Signaling, cell fate, and drug action are all inherently quantitative, stochastic and dynamic phenomena. Predictions for a multitude of contexts and drugs, many of which would not have yet been calibrated for, is facilitated by mechanistic as opposed to empirical models (Birtwistle et al., 2013), so we take a mechanistic chemical kinetics approach.

**Figure 1.**
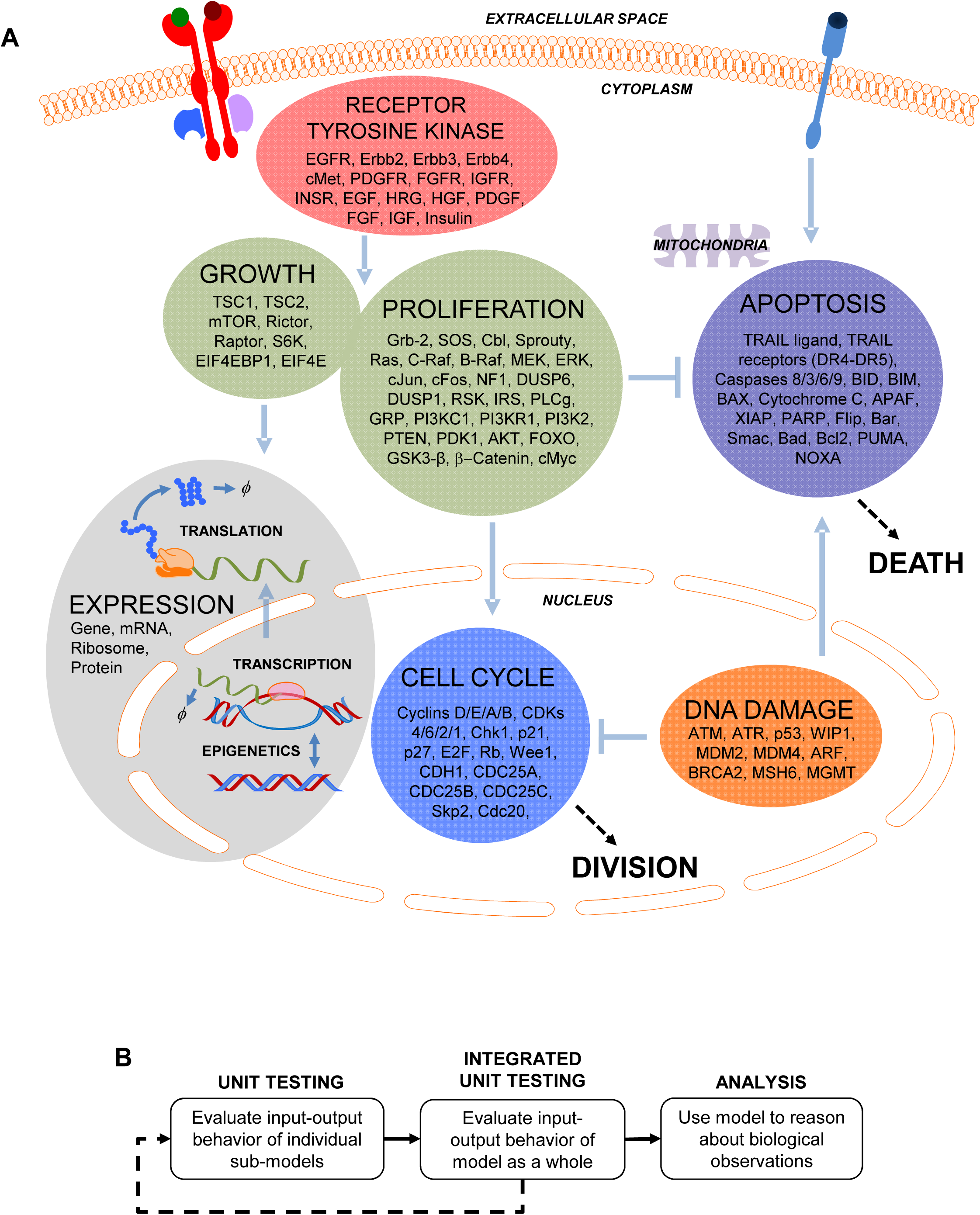
Model overview. A) RTK, proliferation and growth, cell cycle, apoptosis, DNA damage, and gene expression submodels, with genes, compartments and connections indicated. Detailed kinetic mechanisms are in Fig. S1. B) We first evaluate individual submodel behavior (unit testing), followed by integrated unit testing of all submodels together. We then use the model to reason about biological observations.

We present the model as follows (Fig. 1B). In “unit testing” we adapt and develop sub-models for individual pathways (see Supplement) in isolation, requiring that each satisfies a set of experimentally observed biological behavior. We focus on a widely-studied non-transformed cell line, MCF10A (Debnath and Brugge, 2005) to leverage literature data and establish a baseline for “normal” cell behavior before describing transformed contexts with genetic instability and multiple poorly understood mutations. We then connect these sub-models together via established signaling mechanisms to perform “integrated unit testing”, which is iterative. Finally, we analyze the model to reason about emergent signaling mechanisms underlying biological observations.

### Unit Testing—Expression

#### Computationally efficient simulation of stochastic gene expression with expected cell-cell variability

A major driver of cell-to-cell variability in phenotype and drug response is stochastic gene expression (Spencer et al., 2009). We derived a hybrid stochastic-deterministic model that captures random gene switching and mRNA synthesis/degradation, but treats as deterministic protein synthesis/degradation and the associated protein-level biochemistry since they involve much higher molecule number (Fig. 2A and S2A). This expression sub-model produces typical coefficient of variation (CV) of mRNA and protein levels across a simulated population of cells (Fig. 2B).

**Figure 2.**
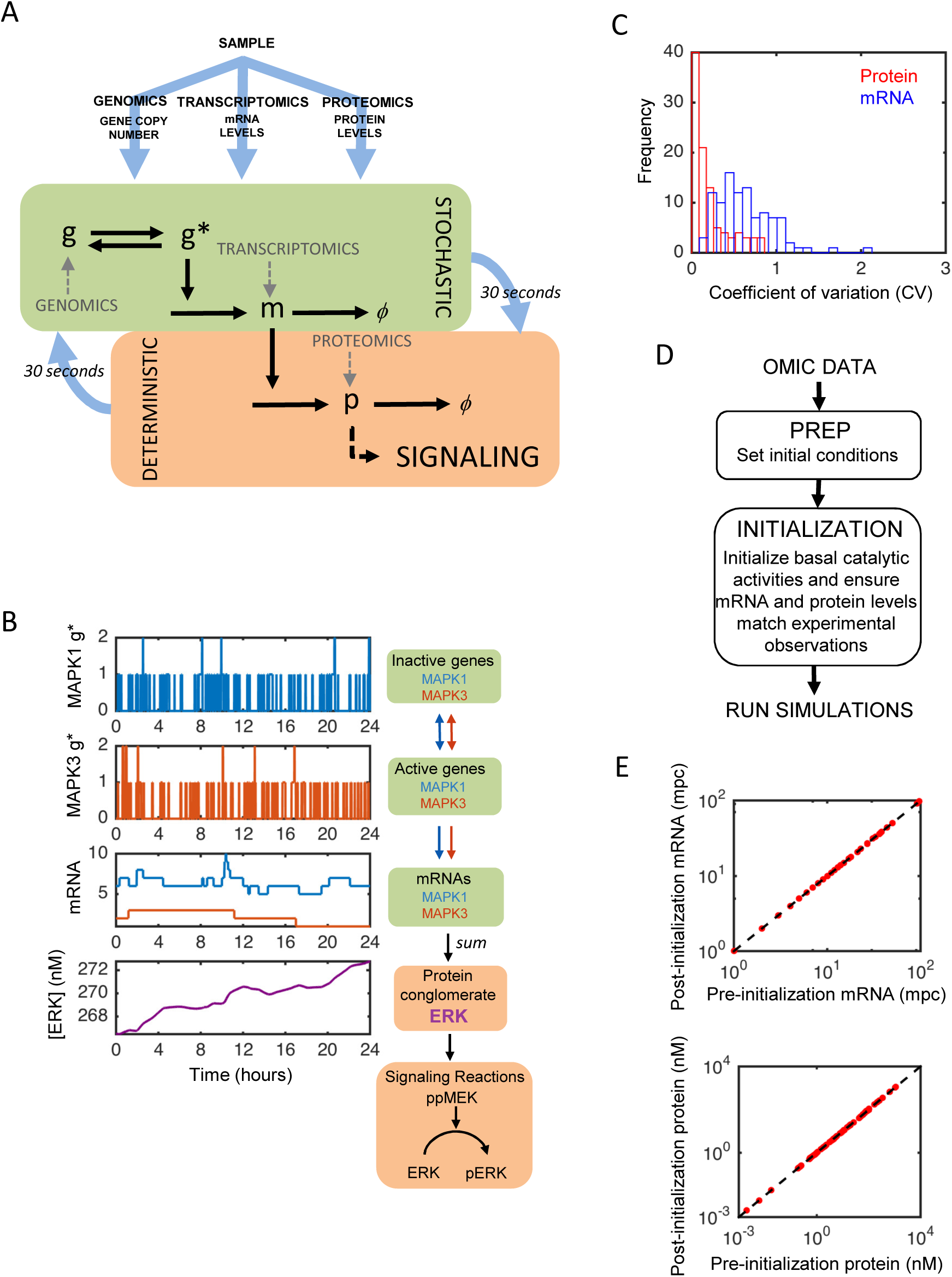
Context tailoring with stochastic gene expression. A) Gene copy number, mRNA levels, and protein levels inform initial conditions and expression submodel parameters. Gene switching, mRNA synthesis, and mRNA degradation are stochastic, whereas all other processes are deterministic, with a 30 second update interval. B) Example simulations for ERK. Two ERK isoform genes (*MAPK1*-blue and *MAPK3*-red) switch between active and inactive states, leading to transcription noise. Transcripts are summed into the signaling conglomerate ERK. C) Coefficient of variation (CV) of protein (red) and mRNA (blue) levels across a population of 100 serum-starved simulated cells. D) Multi-omic data sets initial conditions, then an initialization procedure maintains consistency with experimental observations in the presence of basal signaling activities. E) Simulated mRNA (top) and protein (bottom) levels match experimental data. Dashed line is x=y.

#### Tailoring to omic-data in the MCF10A context

Gene copy numbers in MCF10A (predominantly diploid) have been previously reported (Bessette et al., 2015). We serum-and growth factor-starved MCF10A cells for 24 hours and then quantified molecules / cell of mRNA and protein levels with mRNA-seq and proteomics. These measurements, with prior protein and mRNA stability estimates (see Methods), allow calculation of context- and gene-specific transcription and translation rates (Fig. 2C-D). Many signaling activities consist of groups of genes (e.g. ERK1/2-Fig. 2E); we lump such expressed proteins into relevant “conglomerates” (see Supplement).

#### Control of Translation and Ribosome Dynamics

When cell growth pathways are activated, translation rates (through release of EIF4E) and ribosome synthesis (through p70S6K) increase. We derived a rate law that incorporates the effect of free EIF4E on global translation rates, which cascade into translation rates for each gene product (Methods and Fig. S2B). Ribosome number must be approximately doubled within the course of the cell cycle, which constrains the rate of ribosome synthesis, and also leads to cell growth which we model as increases in volume (Fig. S2C). Simulations reproduce observed positive correlations in cell-to-cell variability for protein pairs (Fig. S2D) (Gaudet et al., 2012).

### Unit Testing—Receptor Tyrosine Kinases

#### Ligand-Receptor Cooperativity

Simulated receptor-ligand binding has reasonable agreement with experimentally reported cooperativity (Fig. 3A). One discrepancy is HRG, showing no cooperativity here but was reported to have negative cooperativity (Hiroshima et al., 2012). HRG binds to ErbB3 and ErbB4 receptors which heterodimerize with ErbB2, all of which have context-specific expression. ErbB2 does not bind ligand and is therefore predicted to make cooperativity less negative for this class of homo and heterodimerizing receptors (including ErbB1/EGFR). Indeed, simulations where receptor expression is better matched make HRG cooperativity more negative (Fig. S3A).

**Figure 3.**
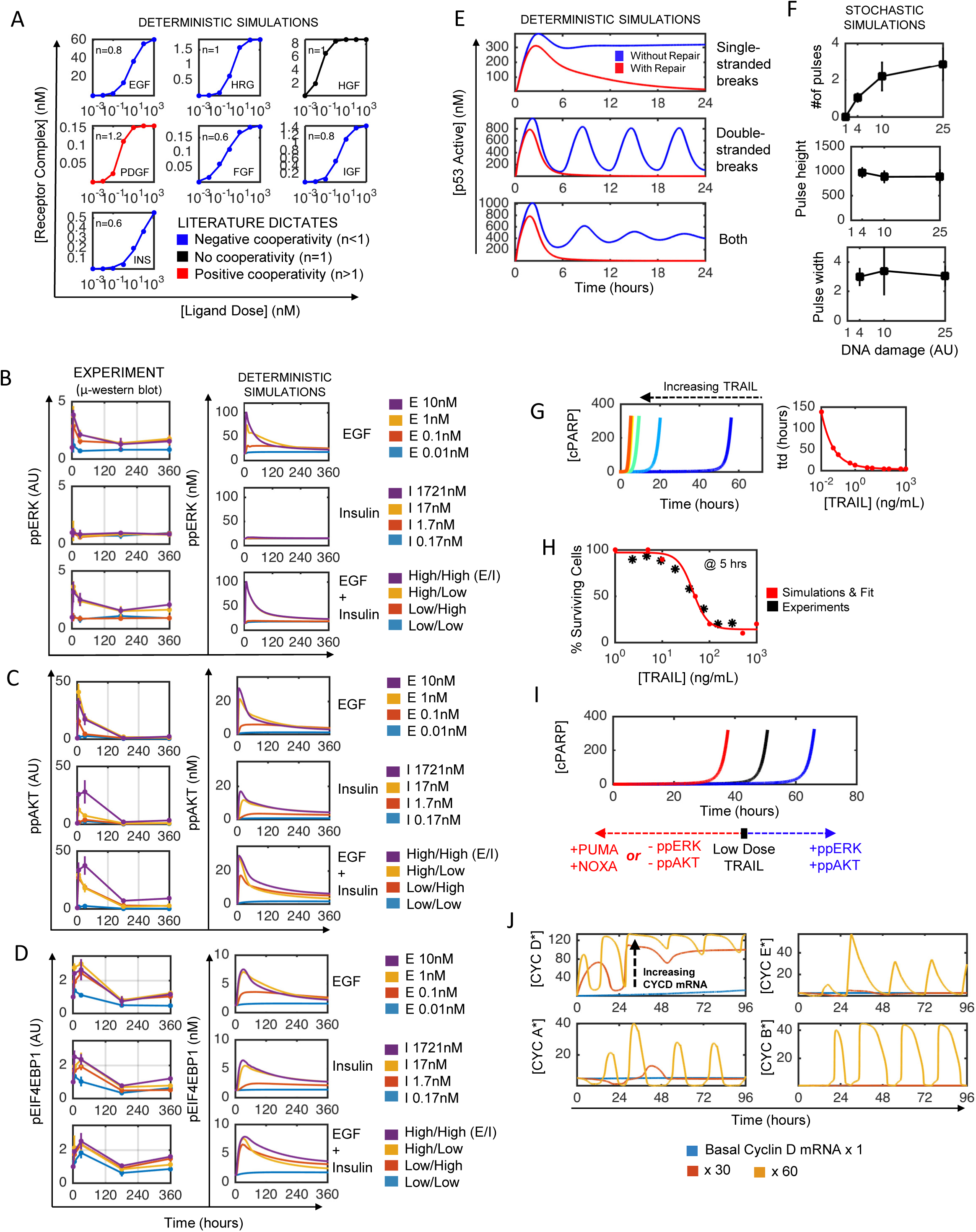
Unit testing. A) Ligand-receptor cooperativity for RTK submodel (dots-deterministic simulations; lines-hill equation fit). B-D) μ-Western blot data (left column) used to train simulated (right column) ERK (B), AKT (C), and mTOR (p4EBP1) (D) signaling dynamics in the proliferation and growth submodel. Serum and growth factor starved MCF10A cells were stimulated with indicated doses of EGF, insulin, and combinations (High/Low EGF 10/0.01 nM, High/Low Insulin 1721/0.17 nM) for 5, 30, 180, or 360 minutes. E) DNA damage-induced p53 responses to single-stranded breaks (top), double-stranded breaks (middle), and both (bottom), with (red) and without (blue) simulated repair. F) Stochastic simulations for how the number of p53 pulses, pulse height (middle) and width (bottom) depend on DNA damage level. G) Increasing amounts of TRAIL (increasingly warmer colors) decreases time to death (ttd—cPARP) exponentially (right panel: red dots-simulation; line-exponential fit). H) Survival after 5 hours TRAIL treatment. Red dots-simulation; red line-sigmoid fit; black dots-flow data. I) Effects of PUMA, NOXA, ppERK and ppAKT on time-to-death in deterministic simulations. J) Simulated cell cycle initiation by increasing cyclin D mRNA levels in serum-starved cells.

#### Receptor Activation and Trafficking Dynamics

Large ligand doses activate (i.e. receptor or adapter phosphorylation) receptor pools within ∼min time scales that are coincident with receptor-specific trafficking dynamics occurring on the order of 5 minutes to 1 hour (Shankaran et al., 2012). Modeled RTKs were consistent with experimental observations for surface vs. internalized receptors and previous models, such as the transient nature of EGF-induced signaling relative to other ligands (Fig. S3B).

### Unit Testing—Proliferation and Growth

#### Receptor Pathway Preferences

We incorporated receptor/adapter affinity and avidity patterns based on prior experimental literature and computational models (see Supplement). For example, ErbB3, IGF1-R and Ins-R all favor PI-3K/AKT signaling heavily as compared to Ras/ERK signaling, which is recapitulated by our model (Fig. S3B). Because our model is tailored to the MCF10A context, certain ligand simulations (FGF, PDGF) do not induce appreciable activation of the ERK or AKT pathways (Fig. S3C), likely due to low receptor expression. This observation is consistent with public data available in LINCS (Fig. S3D) (Niepel et al., 2014), with the exception of IGF, which has greater ERK activation than predicted.

#### Basal Activity Levels

The model describes futile cycles through the Ras and PI-3K pathways (GTP bound Ras and PI(3,4,5)P), and thus accounts for the natural thresholding and negative feedback that prevents such basal activity from influencing phenotype, regulates cellular homeostasis and provides robustness to noise. This causes basal ERK and AKT kinase activity, which constrains phenotype predictions during integrated unit testing (below).

#### Dose and Dynamic Responses

We collected dose-dependent time course data using the microwestern array (meso-scale quantitative western blot) for the ERK (pERK), AKT (pAKT), and mTOR (pEIF4E-BP1) signaling pathways in serum-starved cells treated with to EGF, insulin or the combination (Fig. 3B-D and S4). Due to the varied and dose-dependent pathway preferences of these ligands, we reasoned this dataset would provide constraints on the downstream fluxes. By varying relative fluxes through activated receptors to Ras, PI-3K, ERK, AKT and mTOR, simulations agree reasonably well with this entire cohort of data simultaneously (Fig. 3B-D). These data show the strong influence of Insulin on AKT but not ERK signaling, and also transient EGF-induced signaling as reported in multiple previous studies (e.g. Borisov et al., 2009).

### Unit Testing—DNA Damage

#### DNA repair

We incorporated DNA repair to either single/point damage (MSH6, MGMT) or double (BRCA2) strand breaks (Fig. 3E). We required that levels of damage significant enough to induce p53 response should be repaired within the hours-day time scale.

#### Stochastic regulation of p53 pulsing amplitudes and dynamics

Double strand breaks cause p53 pulses of roughly equal amplitude and timing, but increase in number as DNA damage dose increases, with this number varying across a cell population (Lahav et al., 2004). We specified ultrasensitive activation of ATM and ATR by DNA damage and tuned DNA repair activities (properties hypothesized by (Lahav et al., 2004)) to ensure our model recapitulated these features (Fig. 3F and S3E).

### Unit Testing—Apoptosis

#### Extrinsic pathway dose and dynamic response

The time-to-death of a cell decreases with increasing TRAIL dose (Albeck et al., 2008), which simulations reproduce (Fig. 3G), like the source model. Stochastic simulations match our percent death data for the TRAIL dose response (Fig. 3H), similar to previous work (Flusberg et al., 2003).

#### Survival signaling and intrinsic pathway

DNA damage causes p53-mediated upregulation of pro-apoptotic PUMA and NOXA, but survival signaling activity of the ERK or AKT pathways can counteract this stimulus. The model captures these general trends of PUMA/NOXA and ERK/AKT activities on time-to-death (Fig. 3I).

### Unit Testing—Cell Cycle

#### Expected order and timing of cyclin/cdk complex activities by Cyclin D induction

Simulated Cyclin D transcriptional activation initiates the cell cycle (Fig. 3J). As in the original model, the order and timing of Cyclin expression is maintained (Gérard and Goldbeter, 2009), and these orders correspond to distinct phases of the cell cycle (Fig. 3J). However, stochastic gene expression caused cell cycle deregulation, as found previously (Kar et al., 2009), suggesting as-yet undiscovered mechanisms of cell cycle robustness. We could not rectify this so cell cycle species do not couple to stochastic gene expression.

### Integrated Unit Testing—Model Initialization in a Serum-Starved State

Each sub-model unit is now constrained by a variety of biological observations and data, so we move to integrated unit testing. We do not imply that sub-models are “correct”. Rather, the presented characterization simply increases confidence that sub-models reproduce expected cellular behavior, which of course is predicated upon the considered data and assumptions. We first evaluate integrated behavior in the reference state—serum starvation.

#### Adjusting translation due to protein dispersion

After translation, proteins undergo post-translational modifications (PTM) and/or are recruited into complexes, which can change degradation rates (e.g. GSK-3β destabilizes β-Catenin). Thus, in the integrated model, total protein levels no longer agree with proteomics data. We therefore iteratively adjust effective translation rate constants to retain such agreement (Fig. 2C-D).

#### Cell fate dynamics

Flow cytometry showed that in serum- and growth factor-starved cells state, approximately 10% more cells die every 24 hours (Fig. 4A), and active cell cycling is negligible. These data constrained the effects of proliferation and growth sub-model outputs (primarily ERK and AKT activities) on cyclin D expression (cell cycle) and BAD/BIM (apoptosis), and thus the effects of basal signaling activities on phenotype.

**Figure 4.**
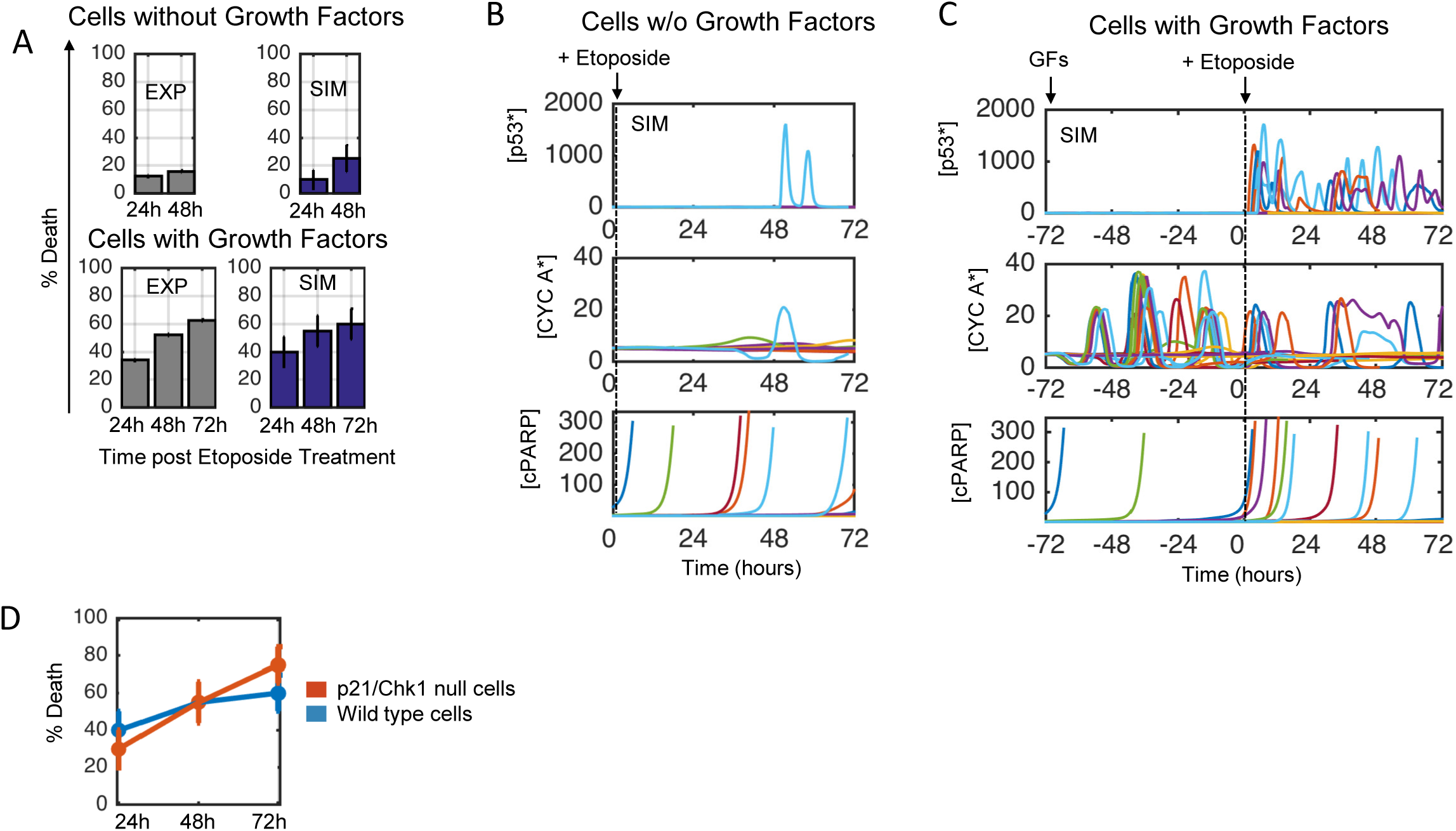
Etoposide response dynamics in dividing and non-dividing cells. A) Experiments (left, grey, flow cytometry) and simulations (right, blue) for MCF10A cells treated with Etoposide (100μM) for 24, 48, and 72 hours in the presence of no growth factors (top) or EGF+Ins (bottom). B-C) Stochastic simulations for (B) serum-starved or (C) EGF+Ins with 72 hr Etoposide treatment. D) Quantification of simulation results from (C) for wild type (blue) or p21 loss cells (red).

### Integrated Unit Testing and Analysis—Cell Cycle-Dependence of Chemotherapy

The chemotherapeutic etoposide induces DNA damage. We treated serum-starved MCF10A cells with etoposide for 24 or 48 hours, but found negligible induction of cell death over control (Fig. 4A-top). This is consistent with etoposide cell death being amplified by DNA replication (Sullivan et al., 1987), which we also observed (EGF+insulin, Fig. 4A-bottom). We postulated a pharmacodynamic model where etoposide-induced DNA damage is strongly dependent on S-phase cyclins (see Supplement) that reproduced the observed cell death results (Fig. 4B-C). Very few simulated cells without growth factors show a p53 response or active cycling (Cyclin A), and therefore very little cell death (cleaved PARP). With EGF and insulin most simulated cells show a robust and sustained p53 pulsing response, which leads to cell cycle arrest (through p21 and Chk1) and cell death.

We noticed that etoposide-induced cell death in the presence of EGF and insulin plateaus over time in experiments and simulations at around 60%, as opposed to increased cell killing over time. We reasoned that this was due to p21/Chk1-induced cycle arrest. In simulations where p53 has limited ability to upregulate p21, etoposide is predicted to cause greater cell death (Fig. 4D). This result is supported by studies showing that cells with mutated p21 are more sensitive to DNA damaging agents (Roninson, 2002; Waldman et al., 1996, 1997). Furthermore, several cancer cell lines from the Genomics of Drug Sensitivity in Cancer Project with p21 mutations exhibit increased sensitivity to etoposide (Garnett et al., 2012). Thus, our model recapitulates the complex dynamic response to a common chemotherapy and predicts features of its context-dependent efficacy.

### Integrated Unit Testing and Analysis—Synergistic Inhibitors for Stochastic Apoptosis

The mechanisms by which oncogene-targeted drugs cause cell death through loss of survival signaling are not as well understood as their cytostatic effects. We measured cell death dynamics (Annexin-V flow cytometry) in response to small molecule inhibitors of ERK and AKT pathways (downstream mediators of many oncogene-targeted drugs) alone or in combination, with serum-starvation or EGF+insulin (Fig. 5A). As before, serum-starvation induces cell death, which is attenuated by EGF+insulin. Inhibition of either ERK or the AKT pathway alone increases cell death 72 hr post treatment, but negligibly at 48 hrs. Combining the two inhibitors revealed synergy in cell death induction at 48 hrs, and much more cell death at 72 hrs. The integrated model captures these cell death dynamics reasonably well (Fig. 5A—far right).

**Figure 5.**
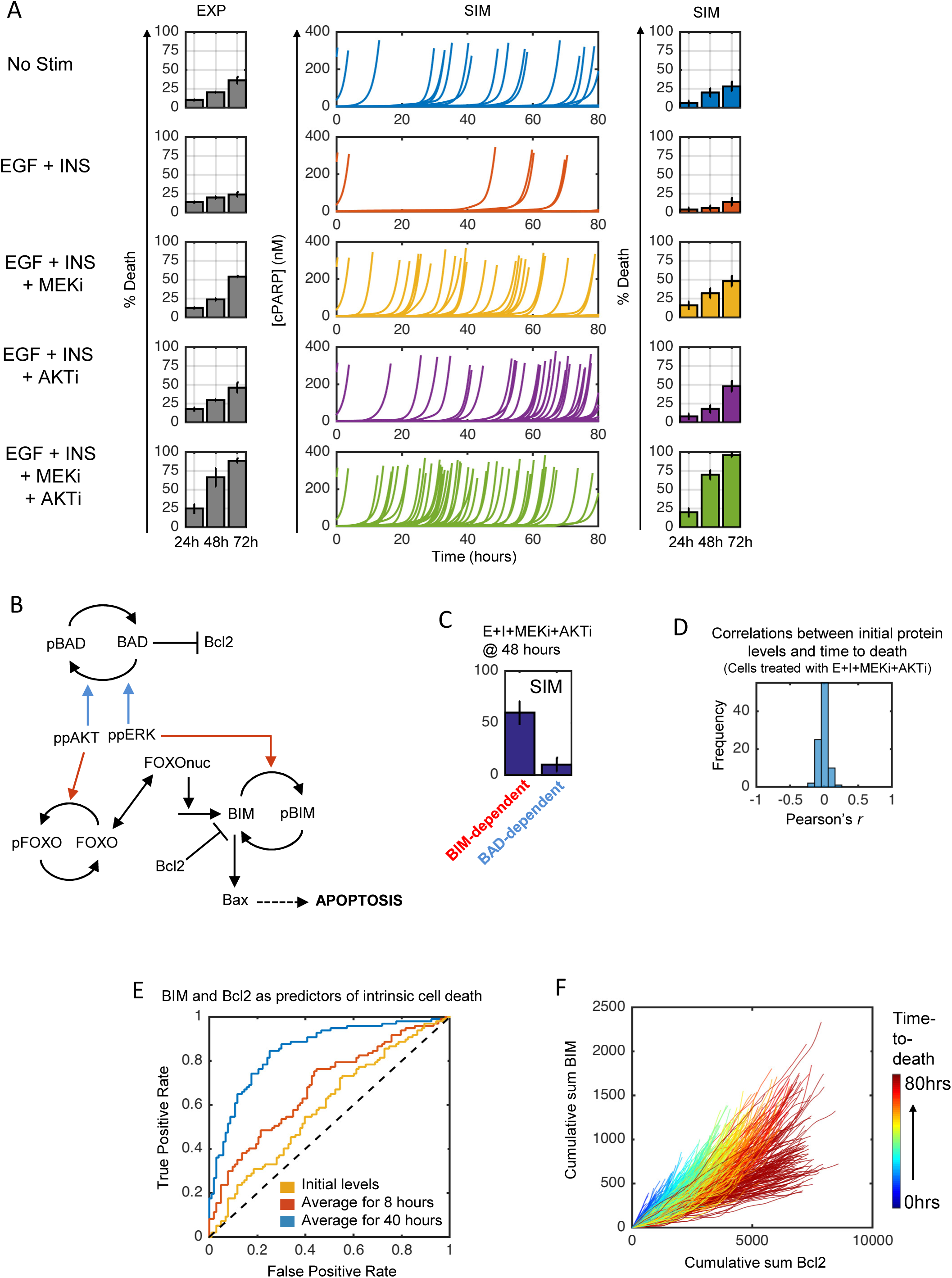
Intracellular regulation of stochastic apoptosis. A) Starved MCF10A cells were treated as indicated (EGF-3.3 nM, insulin-1721 nM, MEKi-10uM, AKTi-10uM) and assayed for cell death by flow cytometry. Single cell simulations (middle, each line) show time to death (sharp cleaved PARP increase). Colors represent treatment condition. B) Modeled ppERK and ppAKT regulation of apoptosis through BAD (blue arrows) and BIM (red arrows). C) Simulations where BAD-dependent (blue arrows in B) or BIM-dependent (red arrows in B) mechanisms were switched off and % death calculated. D) Correlation of initial protein levels with time-to-death for 400 simulated cells treated with EGF+ins+MEKi+AKTi. E) ROC curves for predicting cell death from time-averaged BIM and Bcl2 levels based on 400 simulated cells (200 training/200 validation) treated as in (D) (blue-AUC=0.85; red-AUC=0.68; yellow-AUC=0.59). F) Cumulative sum of BIM and Bcl2 levels for simulated cells before death for all conditions in (A). Color shows time to death.

We used the integrated model to reason about what mechanisms may underlie the synergy between and overall impacts of ERK and AKT inhibition for cell death. Both ERK and AKT can phosphorylate and inactivate the pro-apoptotic function of BAD, ERK can phosphorylate BIM causing similar inhibition of pro-apoptotic function, and AKT-mediated activation of FOXO upregulates BIM levels (Fig. 5B). Simulations where either BAD-(blue arrows) or BIM-dependent (red arrows) mechanisms were computationally knocked-out predicted BIM to account for most ERK and AKT inhibition effects (Fig. 5C). Independent experimental data corroborate this prediction; MCF10A cell detachment-induced death, which depends on such survival signaling mechanisms, has been observed to be predominantly controlled by BIM rather than BAD (Reginato et al., 2003). The model suggests several contributing mechanisms. One reason is expression levels; BAD is lowly expressed compared to BCL2 family proteins (13,709 vs. 61,003 molecules per cell), and thus is unlikely to exert significant control of cell death. Another is the cascade network structure for BIM regulation. If only AKT is inhibited, phosphorylation by ERK blocks the accumulation of active BIM. If only ERK is inhibited, the amount of BIM in the cell, even if completely unphosphorylated, is largely insufficient to drive cell death in the absence of further strong death signals. However, when both pathways are inhibited, the production and accumulation of active BIM proceeds uninhibited, thus creating a potent apoptotic signal. Furthermore, the inhibition of these activities must be coordinated over relatively long periods of time, with duration of AKT inhibition having enough overlap with that of ERK to cause both the accumulation and activation of BIM.

Why do some cells die early and others die late following ERK and AKT inhibition? We used simulation data to gain insight into this question. First, we asked whether initial total protein levels (sum across all forms of a protein) were correlated with time-to-death after ERK and AKT inhibition. In line with conclusions from prior studies focused on extrinsic death pathways (Spencer et al., 2009), time-to-death was not significantly correlated with the initial levels of any single protein (Fig. 5D). This suggests that stochastic processes post-inhibitor treatment largely dictate variable cell death fate. We performed lasso regression using time-averaged protein levels as predictors of early or late death (pre/post 40 hours) over 400 individual cell simulations. This analysis identified time-averaged BIM and BCL2 as the top explanatory variables. Next, we trained a support vector machine (200 cells for training, 200 for validation) and found that average BIM and BCL2 levels over a 40-hour time course (or as long as a cell lived) are highly predictive for death timing (Fig. 5E). Initial levels, or even those averaged over the first 8 hours post-stimulus, were moderately predictive at best. As one might expect based on the respective pro- and anti-apoptotic functions of BIM and BCL2, respectively, stochastic expression time courses that tend towards high BCL2 and low BIM are the most protective (Fig. 5F). Interestingly, these predictors of stochastic death phenomena were not equally applicable to different death stimuli such as the extrinsic ligand TRAIL (Fig. S5A), thought to depend more on stochastic activation dynamics of Caspase 8 (Spencer et al., 2009), which highlights the treatment-specificity of stochastic cell fate determinants. We conclude that intrinsic death of individual cells induced by inhibition of survival signals is likely to be inherently unpredictable prior to treatment.

### Integrated Unit Testing and Analysis—Synergistic Mitogens for Stochastic Cell Cycle Entry

EGF and insulin regulate MCF10A proliferation. We measured (BrdU incorporation/flow cytometry) how these two ligands influence cell cycle progression of serum-starved MCF10A cells and how the ERK and AKT pathways were involved (Fig. 6A). Insulin induces negligible cell cycle entry on its own but synergizes with EGF, which is also seen on the level of cyclin D expression (Fig. 6B, left). The ERK and AKT pathways are both essential to drive EGF+insulin-induced cell cycle progression (Fig. 6A). After tuning the dependencies of transcriptional processes downstream of the ERK and AKT pathways, the integrated model reproduces these data (Fig. 6B-C).

**Figure 6.**
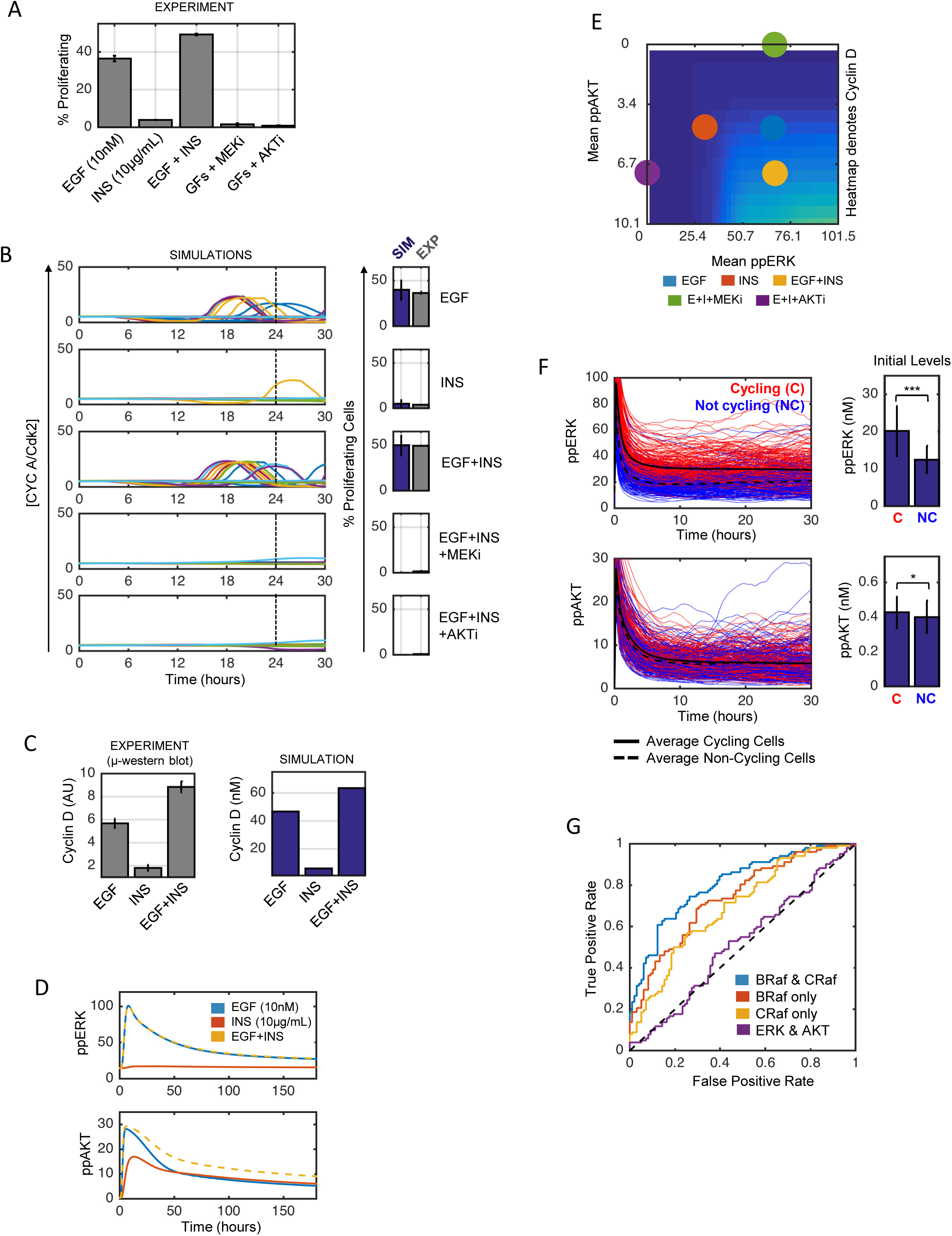
Regulation of stochastic proliferation. A) Starved MCF10A cells were treated as indicated for 24 hours and then assayed for cell cycle progression (flow cytometry). Percent proliferating cells are S-phase plus G2/M minus that in serum-starved controls (<∼5%). B) Model fit. C) Measured (left) and simulated (right) cyclin D levels 6 hours post stimulation of starved cells. D) Deterministic simulations of ppERK (top) and ppAKT (bottom) response of starved cells to indicated growth factors. E) Simulated relationships between time-averaged ppERK and ppAKT with cyclin D levels (heatmap colors). Colored circles indicate values for different treatments. F) 400 simulated cells treated with EGF+Ins were cycling (red; divided before 30 hours) or non-cycling (blue; did not divide before 30 hours). Right: Initial levels of ppERK or ppAKT in the same simulations. G) ROC curves predicting proliferation from 400 simulated cells (200 training/200 validation) following EGF+ins from initial BRaf and CRaf levels (blue-AUC=0.81; red-AUC=0.74; yellow-AUC=0.69; purple-AUC=0.51).

What underlies the synergy between EGF and insulin with regards to cell cycle progression? Prior work has suggested that the dynamics of the ERK pathway can determine cell proliferation fate (Nakakuki et al., 2010). However, data and simulations suggest that ERK pathway activation dynamics are essentially identical for EGF or EGF+insulin (Fig. 6D). Insulin is a very strong activator of AKT but not ERK signaling (Fig. 6D). When cells are treated with EGF+insulin, there are negligible differences in acute (<30 min) AKT activation dynamics, and significant differences only develop over long (hrs) time scales. Thus, the mitogenic synergy between EGF and insulin seems associated with prolonged AKT pathway activity over long time scales (Fig. S6). Simulations suggest that increasing mitogen-induced cyclin D expression cannot be accomplished by only increasing activity of one pathway, but rather roughly equal amounts of both time-integrated ERK and AKT activity are needed to drive more robust mitogenic responses (Fig. 6E). This interpretation cannot rule out coincident factors such as increased insulin-induced glucose uptake, but does suggest that the coordinated activation of both ERK and AKT for several hours post-mitogen treatment seem important to drive cell cycle entry, rather than acute activation dynamics following mitogenic stimulus.

These observations suggested to us that stochastic differences from cell-to-cell in AKT activity dynamics might be predictive for EGF+insulin-induced cell cycle entry. Surprisingly, in simulation results, neither the initial AKT activity nor dynamics post-stimulus contained significant discriminatory information related to subsequent cell cycle entry decisions of individual simulated cells (Fig. 6F). Rather, the ERK activity dynamics were much more predictive of subsequent stochastic cell cycle entry (Fig. 6F). This is consistent with prior live-cell imaging work that showed a strong correlation between time-integrated ERK activity and resultant S-phase entry decisions (Albeck et al., 2013), albeit in a setting of much higher cell confluence than our experiments here entail and with additional mitogenic factors present (hydrocortisone and cholera toxin). Why would AKT activity dynamics control synergy between EGF and insulin but not be predictive of stochastic cell fate? AP1 and cMyc control cyclin D expression in this model. AP1 (cFos-cJun) activity has a sharp response to time-integrated ERK activity, relative to the less sharp dependence of cMyc expression on AKT activity. This sharpness difference is due to positive autoregulation of cJun expression by AP1 (Angel et al., 1988), and is likely also influenced by the already high expression of cMyc in serum-starved MCF10A (47,902 molecules/cell, compared to 0 and 3,101 for cFos and cJun), which is one of the few MCF10A alterations (they also have CDKN2A/ARF loss, which is captured in our gene copy number data) (Kadota et al., 2010).

Could stochastic cell cycle entry be predicted by protein abundance fluctuations prior to EGF+insulin treatment? We again applied lasso regression using initial conditions for total protein levels in individual simulated cells to identify explanatory variables for cell cycle entry, and then trained a support vector machine classifier. In contrast to predicting death in individual cells, here we found that initial CRaf and BRaf levels were fairly predictive of whether a cell would center the cell cycle by 24 hrs post EGF+insulin stimulus (Fig. 6G). Initial total ERK and AKT (active + inactive) levels were not predictive at all (Fig. 6G). We conclude that so long as AKT activity is prolonged, ERK activity fluctuations likely control stochastic cell proliferation fate following acute growth factor stimulus, which are to predominantly set by initial Raf protein levels.

### Analysis—Predicting Initial Stages of Transformation Fitness

The model contains multiple oncogenes and tumor suppressors, and is calibrated to a non-transformed epithelial cell state. Does our model have predictive ability for transformation potential? To test this, we systematically altered expression of each RTK, proliferation and growth sub-model gene (10-fold up for oncogenes, 10-fold down for tumor suppressors), simulated proliferation response to EGF+insulin (emulate a growth factor-containing microenvironment), and ranked nodes by this metric (Fig. 7A). Several expected high-ranking results are observed, such as Ras, Raf, ErbB2/HER2 and EGFR, those with well-established transforming viral analogs such as cFos, cJun and cCbl, and more recently recognized translation control with EIF4E and 4E-BP1 (Musa et al., 2016). Gene products along the PI3K axis (PIK3Cα, PTEN, β-Catenin, cMyc) are lower ranked, which seems counter-intuitive since these are also frequent cancer drivers. However, MCF10A cells, while non-transformed, are immortal and fast growing in culture indefinitely, and one of the reasons (as stated above) is cMyc overexpression, downstream biochemically of PI3K/AKT and is a resistance mutation to PI3K therapy that tends to be mutually exclusive (Dey et al., 2015; Miller, 2012; Moumen et al., 2013). Thus, we interpret these results as being consistent with the MCF10A context being naturally less reliant on these PI3K axis gene products to drive increased proliferation.

**Figure 7.**
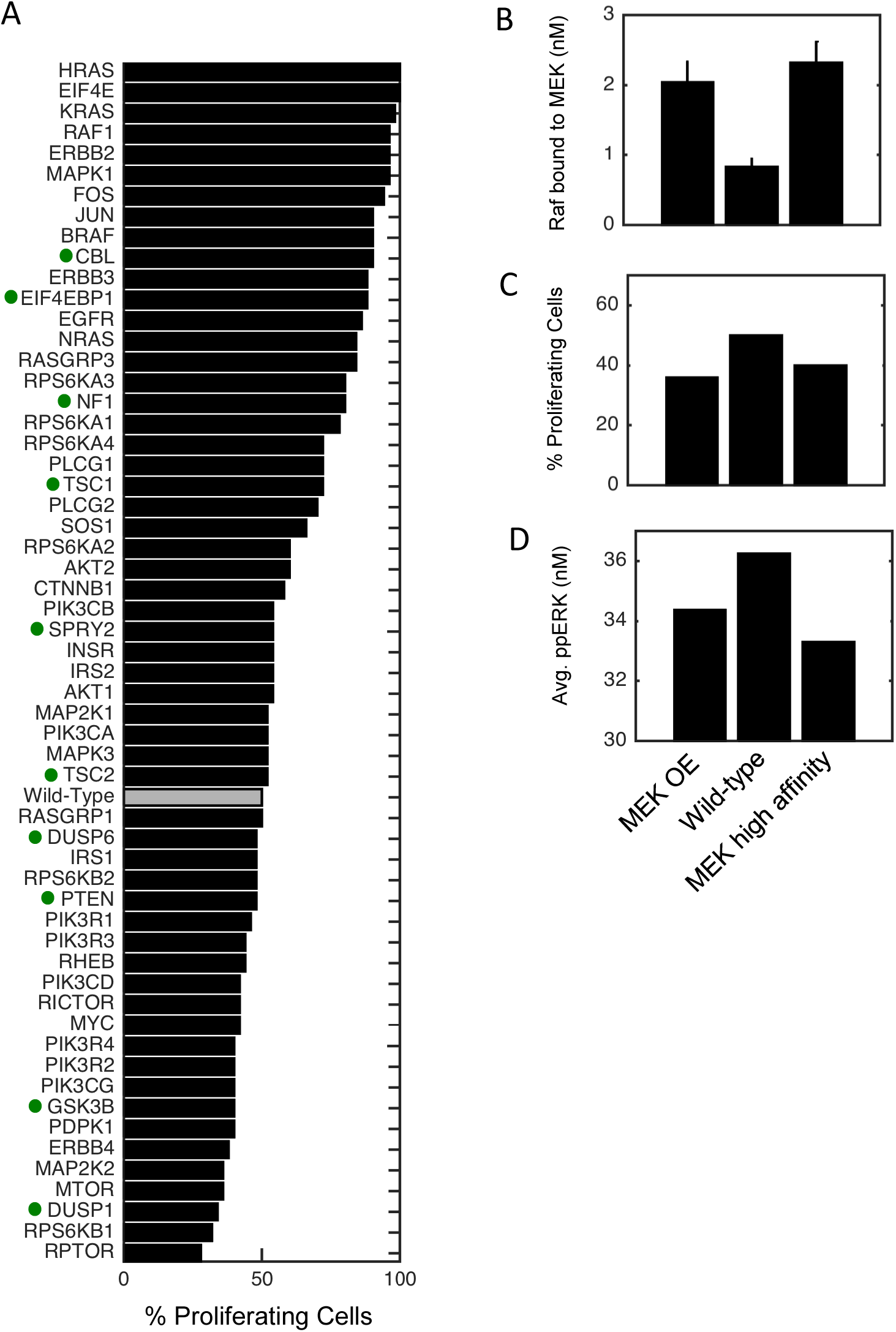
Simulated Transformation. A) The mRNA level of each gene is altered 10-fold and EGF+Ins response of starved cells for 24 hrs is simulated. Gray bar denotes wild-type and dots denote tumor suppressors (10-fold reduction). B-D) MAP2K2 overexpression (left), wild-type (middle), and MEK-Raf high affinity (right, unphosphorylated MEK has 10-fold higher affinity for all Raf species) were simulated as above and either the level of Raf bound to MEK at t=0 (B), % proliferating cells (C) or time-averaged ppERK levels (D) were calculated.

One paradoxical result is the low ranking of MEK (MAP2K1/2), despite being functionally surrounded by highly ranked genes. This result is actually consistent with widespread clinical data that seldom report MEK alterations (∼<5% TCGA pan-cancer in cBioPortal) as compared to Ras/Raf alterations, the explanation for which remains unclear. What mechanisms might explain MEK’s poor simulated transforming potential? In simulations, free C/B-Raf—those that are available to transmit signals from active Ras—are reduced, and most of this difference ends up sequestered by MEK itself, which seems to reduce active ERK (Fig. 7B-D). Simulations where MEK has artificially high affinity for MEK phenocopy this result. We interpret this as a consequence of stoichiometry and expression levels, since both C-Raf and B-Raf are relatively lowly expressed (28,849 mpc and 2,825 mpc, respectively) as compared to MEK (MAP2K1-147,250 mpc; MAP2K2-377,540 mpc), so although this sequestration effect might be relatively small, it can account for a non-negligible proportion of the total Raf pool. Thus, the prediction from these simulations is that MEK typically has poor transforming potential because increasing its levels it tends to sequester Raf from increased signaling as opposed to allowing for increased signal transmission. This of course does not explain observations related to activating mutations, which convolve protein structural robustness, and also potentially genome organization factors. Such a mechanism would not be possible to uncover from genomics or network model-based approaches because it is inherently based in quantitative biochemistry that our model captures from its formulation.

## DISCUSSION

We developed the first mechanistic pan-cancer driver network model of cell proliferation and death regulation that is tailorable to biological context defined by multiomics data. The model describes fractional cell kill and partial cytostasis in response to treatment with a variety of anti-cancer drugs, conditioned upon an expression and microenvironmental context. We first developed sets of unit test operations that build confidence in individual submodels, and then evaluated the ability of the model to reproduce cell biological behavior as an integrated whole, which we subsequently analyzed to gain insight into biochemical mechanisms of stochastic cell proliferation, death and transformation.

While this model is complex and larger than any previous such signaling model, some may argue it still does not account for enough biology (e.g. post-translational Myc regulation and varied transcriptional targets), while yet others may argue that it is too large to be useful. Symmetric critiques often reflect striking an appropriate balance, but both views have validity. We have only focused on a single non-transformed cell line with no mutations. Impeding future work here is that the most mutations are not functionally well-understood. We also do not consider tumor metabolism or many signaling pathways important outside pan-cancer scope. Rule-based modeling (Faeder et al., 2009) and algorithmic model building (Lopez et al., 2013) could help modelers capture more biology. Open and FAIR (Wilkinson et al., 2016) data that is well-annotated and available by computational query (e.g. LINCS consortium) will be needed to drive such modeling forward.

We certainly do not claim that our present model is unique, neither in terms of structure nor parameterization. These classes of models are well-known to exhibit parametric non-identifiability but yet are still able to make valid and precise biological predictions (Gutenkunst et al., 2007). The fact that these models stand on the shoulders of decades of cell and molecular biology research and are formulated to the extent possible based on physicochemical principles facilitates predictive potential despite the inherent uncertainty. Increased ability to algorithmically compare model structure and predictions to a wide net of publicly available experimental data will help reduce model uncertainty. Regardless, model complexity is driven by the questions and data at hand, and here the ultimate goal of cancer precision medicine coupled with omics data for tailoring gives a rational basis for the model size and complexity.

To study stochastic cell fate with our model we combined penalized regression (identify predictive features) with machine learning (relate predictive features to cell fates). Like prior work (Spencer et al., 2009), we found that ERK+AKT inhibitor induced apoptosis fate was not predictable from initial protein levels. Rather, the couplet of time-averaged total BIM and BCL2 levels following death stimulus were highly predictive. Thus, apoptosis seems inherently unpredictable at the time of the death stimulus, and rather evolves stochastically over the treatment time course.

In contrast to the inherent unpredictability of apoptosis based on initial protein levels, S-phase entry in simulations was predictable from initial C-Raf and B-Raf levels. Why is this the case for S-phase entry and not apoptosis? There are several potential factors in the model. First is the relatively low expression levels of both Raf isoforms, giving larger expression noise. Second is the bottleneck role of Raf in propagating signals from RTKs to S-phase entry, coupled with enzymatic signal amplification as opposed to stoichiometric titration as in apoptosis signaling. Third is the predominant stochastic control of S-phase entry by ERK signaling dynamics, as opposed to AKT signaling dynamics. This may be due in part to the downstream positive feedback of cJun expression (which amplifies noise).

Our model can naturally account for drug dose, promiscuity, scheduling, kinetics and pharmacodynamics. These hallmarks of pharmacology are as yet difficult to rationalize with genomic approaches. One potential application of ours and similar models is improving upon genomic-based approaches to cancer precision medicine. For example, patient transcriptomic data could be used to tailor the model as described here, giving an *in silico* test bed to explore a plethora of single or combination therapy options, including dosing and timing. Given advances in patient-derived experimental systems from biopsies (Liu et al., 2012), perturbation data can be used to further refine patient-specific mechanistic models and test drug predictions prior to clinical administration. Such predictions would have engrained within important properties of clinical response such as the promiscuity of many targeted kinase inhibitors (Davis et al., 2011) and dynamic feedback mechanisms in cancer cells important for resistance (Chandarlapaty et al., 2011). Moreover, given inference of subclonal evolution patterns of a patient’s tumor (Landau et al., 2013), multiple instantiations of the model could be generated to find a cross-section of therapies, or perhaps even order of therapies, that could most efficiently control an (epi)genomically heterogeneous tumor.

Alternatively, virtual clinical trials could be conducted for a particular drug or combination, given a cohort of patients defined by multi-omics data (e.g. transcriptomics). Simulation results could stratify patients for inclusion or exclusion in trials—an exceedingly important task given the niche efficacy of targeted drugs from single genetic biomarkers. Further simulations could optimize dosing and timing strategies. With now hundreds of FDA-approved anti-cancer therapeutics (including dozens of targeted drugs), and the realization that combination therapy is almost always required for clinical response, computational approaches such as this that help prioritize drugs, their doses, and their scheduling for particular patients will become increasingly important in drug development and personalized oncology.

## AUTHOR CONTRIBUTIONS

MRB conceived of and supervised the work. MB created the model and performed all simulations and flow cytometry experiments. MRB and MB wrote the paper. AMB and RJK performed microwestern experiments. LCS collected TRAIL dose response data. ADS processed cells for mRNA-seq and proteomics. MSD, EAR, AT and AM helped with model building and analysis.

## ACKNOWLEDGEMENTS

MRB acknowledges funding from Mount Sinai, the NIH Grants P50GM071558 (Systems Biology Center New York), R01GM104184 and U54HG008098 (LINCS Center), and an IBM faculty award. MB and ADS were supported by a NIGMS-funded Integrated Pharmacological Sciences Training Program grant (T32GM062754). We thank Jerry Chipuk for helpful discussion, Mohit Jain and Hong Li for proteomics services (funded in part by NIH grant NS046593, Rutgers Neuroproteomics Core Facility), and Evren Azeloglu, Bin (Tina) Hu, Gomathi Jayaraman and Yuguang Xiong for help with mRNA sequencing.

## Supplementary Methods

### CONTACT FOR REAGENT AND RESOURCE SHARING

Further information and requests for resources and reagents should be directed to and will be fulfilled by the corresponding author, Marc Birtwistle.

### EXPERIMENTAL MODEL AND SUBJECT DETAILS

#### Cell culture

MCF10A cells were cultured in DMEM/F12 (Gibco; Cat: 1133032) medium supplemented with 5% (v/v) horse serum (Gibco; Cat: 16050-122), 20ng/mL EGF (PeproTech, Cat: AF-100-15), 0.5 mg/mL hydrocortisone (Sigma, Cat: H-0888), 10μg/mL insulin (Sigma, Cat: I-1882), 100ng/mL cholera toxin (Sigma, Cat: C-8052), 2mM L-Glutamine (Corning; Cat: 25-005-CI) and 1X (10,000 I.U./mL / 10,000 ug/mL) penicillin/streptomycin (Corning; Cat: 30-002-CI). Cells were cultured at 37°C in 5% CO_2_ in a humidified incubator, and passaged every 2-3 days with 0.25% trypsin (Corning; Cat: 25-053-CI) to maintain subconfluency. Cells originate from a female host, were a gift (Gordon Mills and Yoko Irie) and we did not independently authenticate the cells. Starvation medium was DMEM/F12 medium supplemented with 2mM L-Glutamine and 1X penicillin/streptomycin.

### METHOD DETAILS

#### Experimental Methods

##### RNA sequencing and analysis

MCF10A cells were seeded at ∼30% density (1 million cells / 60mm diameter dish, one dish per biological replicate), and incubated overnight in full growth medium. The next morning, the full growth medium was aspirated, cells were washed once with PBS, and then placed in starvation medium for 24 hours. Total RNA was extracted using TRIzol (Life Technologies, Cat: 15596018) per manufacturer instructions (detailed SOP at www.dtoxs.org; DToxS SOP A – 1.0: Total RNA Isolation). RNA sequencing and analysis was performed as previously described (Xiong et al., 2017). We prepared RNA-seq libraries by adapting a single-cell RNA-seq technique—Single Cell RNA Barcoding and Sequencing method (SCRB-seq; Islam et al., 2014; Soumillon et al., 2014)—to extract total RNA from cells (detailed SOP at www.dtoxs.org; DToxS SOP A – 6.0: High-throughput mRNA Seq Library Construction for 3’ Digital Gene Expression (DGE)). This method involves converting each Poly(A)+ mRNA molecule to cDNA labeled with universal adaptors, sample-specific barcodes, and unique molecular identifiers (UMIs) on every transcript molecule using a template-switching reverse transcriptase. cDNA was amplified and prepared for multiplexed sequencing, enriching for 3’ ends preserving strand information by using a modified transposon-based fragmentation approach. Libraries were sequenced using the Illumina HiSeq 2500 platform (detailed SOP at www.dtoxs.org; DToxS SOP A – 7.0: Sequencing 3’-end Broad-Prepared mRNA Libraries). Analysis was performed using a custom Python script (detailed SOP at www.dtoxs.org; DToxS SOP CO – 3.1: Generation of Transcript Read Counts). In brief, sequenced reads were aligned to a 3’-focused hg19 reference using BWA, and counts of reads that aligned to one or more than one gene with high confidence were calculated. The UMI count is defined as the number of distinct unique molecular identifiers (UMI) sequences embedded in those aligned reads and is well-known to alleviate PCR bias that can distort linearity in quantification. Biological triplicates were performed, but one sample failed to be sequenced, leaving biological duplicates.

##### Proteomics and analysis

MCF10A cells were seeded at ∼30% density (5 million cells / 150mm diameter dish, 2 dishes pooled per biological replicate) and incubated overnight in full growth medium. The next morning, the full growth media was aspirated, cells were washed once with PBS, and then placed in starvation medium for 24 hours. Cells were trypsinized for 10 minutes, resuspended in 1X PBS, and spun down at 500g for 5 minutes. Cell pellets were then snap frozen in liquid nitrogen and shipped to the Advanced Proteomics Center at New Jersey School of Medicine to perform mass spectrometry. The cell pellet was subjected to urea lysis buffer composed of 8 M urea, 100 mM TEAB, 1X protease inhibitors (complete, EDTA-free Protease Inhibitor Cocktail from Sigma, prepared according to manufacturer instructions) and 1X phosphatase inhibitors (PhosSTOP from Sigma, prepared according to manufacturer instructions). Protein (50.0 μg total) was resolved by one dimensional SDS-PAGE. The gel lane was divided into 24 equal size bands, followed by in-gel digestion with trypsin performed as in (Shevchenko et al., 2006). Peptides were fractionated by reverse phase chromatography and analyzed directly by LC/MS/MS on a Q Exactive Orbitrap mass spectrometer (funded in part by NIH grant NS046593, for the support of the Rutgers Neuroproteomics Core Facility). We used Maxquant to analyze the data and calculate iBAQ scores, using a human UniProt protein database as the reference proteome. Final proteins were identified with 1.0 % false discovery rate (FDR) at both protein and peptide levels. This resulted in ∼10k unique proteins. Biological triplicates were performed.

##### Calculation of mRNA and Protein Molecules Per Cell

Importantly, the units of the data acquired from the RNA-seq (UMI count; (Islam et al., 2014)) and mass spectrometry (iBAQ scores; (Schwanhausser et al., 2011)) technologies are linearly proportional to the number of mRNA or protein molecules in the original sample, respectively. To convert from these measurement units to molecules per cell, we first quantile normalized the three biological replicates using the MATLAB function, `quantilenorm`, with default settings. The quantile-normalized values were then converted from their original unit of measure to units of molecules per cell with the following logic.

Consider measurement of mRNA or protein *i*, *A*_*i*_ (in UMI counts or iBAQ score), and the corresponding absolute copy number per cell that gives rise to this measurement, *X*_*i*_. Due to the linearity of the measurement modality, we have:

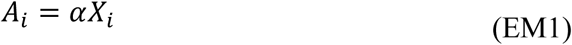

Summing over all species *n* in the cell gives:

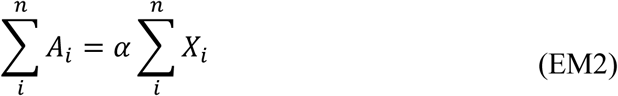

Dividing by these sums then yields the following proportionality to convert the measurement to absolute copy number per cell.

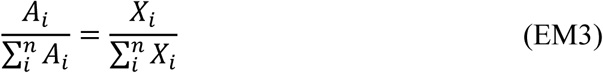

Using these equations requires an estimate for the total number of for mRNA and protein molecules in a cell. The total number of mRNAs in a typical mammalian cell was estimated from three sources. The first was taken from (Schwanhäusser et al., 2011a) by summing all mRNA quantities found in their search (173,921 molecules per cell). The second was taken from the Molecular Biology of the Cell textbook (Alberts, B. Johnson, A. Lewis, J. Raff, M. Roberts, K. Walter, 2008), which gave a value of 360,000 molecules/cell. The third was calculated by converting the total picogram amount of RNA in a typical mammalian cell, estimated from one source to be 26pg (Brandhorst and McConkey, 1974), to molecules per cell given estimates that ∼3% of total RNA is mRNA (Milo et al., 2009), that there are an average of 2000 nucleotides in a single mRNA transcript (Milo et al., 2009), and that the average molecular weight of a single nucleotide is 339.5 g/mol (Milo et al., 2009); using this information we calculated a total 691,891 molecules per cell. We took the average of these three numbers and approximated the result to 400,000 molecules per cell. To approximate the total number of proteins in a typical mammalian cell we summed the total number of protein molecules found by Schwanhausser et al. (2011), which amounted to 3.08 × 10^9^ protein molecules per cell. Importantly, our method for converting to molecules per cell resulted in reasonable agreement with data from Schwanhausser et al. When comparing the numbers between our dataset (“ours”) and the Schwanhausser dataset (“S”) for a handful of constitutively expressed proteins that typically do not significantly change across cell types, we find most of them to be within the same order of magnitude: MAP2K1 (ours: 147250; S: 284958), MAP2K2 (ours: 377540; S: 327073), MAPK1 (ours: 746463; S: 1400595), MAPK3 (ours: 76314; S: 195819), AKT1 (ours: 74762; S: 37060), and AKT2 (ours: 81476; S: 4325).

##### Micro-western experiments and analysis

MCF10A cells were seeded at ∼30% confluence in 6-well plates (150,000 cells / well), incubated overnight in full growth medium. The next morning, the full growth medium was aspirated, cells were washed once with PBS, and then placed in starvation medium for 24 hours. Cells were treated with combinations of EGF and Insulin at various doses for 5 minutes, 30 minutes, 3 hours, or 6 hours. Growth medium was aspirated fully and cells were lysed on ice with a custom-made lysis buffer tailored for micro-western applications composed of 240 mM Tris-Acetate, 1% SDS w/v, 0.5% glycerol v/v, 5mM EDTA, water, and various protease and phosphatase inhibitors—aprotinin (1μg/mL), leupeptin (1μg/mL), pepstatin (1μg/mL), β-glycerophosphate (10mM), sodium orthovanadate (1mM), and dithiothreitol (DTT; 50mM). Reagent sources are listed in our SOPs (www.birtwistlelab.com/protocols). Samples were boiled at 95 degrees for 5 minutes, sonicated using a VialTweeter (Hielscher), then concentrated ∼5 to 10-fold using Amicon Ultra centrifugal filters (Merck Millipore Ltd). Total protein content was assayed via the Pierce 660 assay per the manufacturer’s instructions. Samples were printed in 24 “blots” onto a single 9.5% poly-acrylamide gel using a GeSim Piezoelectric Nanoplotter, followed by horizontal electrophoresis and wet transfer to a nitrocellulose membrane. The membrane was blocked with Odyssey Blocking Buffer (TBS; LICOR) then put into a 24-well gasket system that corresponds to the original lysate printing and horizontal running pattern. This gasket system allows different primary antibodies to be probed in each well. We probed using the following antibodies from Cell Signaling: pERK (Cat: 4370; 1:1000), pAKT (Cat: 4060; 1:2000), pEIF4E-BP1 (Cat: 2855; 1:500), cyclin D (Cat: 2978; 1:500) and α-Tubulin (Cat: 3873; 1:2000). We then incubated the entire gel with two secondary antibodies from LiCor: IR dye 800CW goat anti-rabbit (green; Cat: 926-32211) and IR dye 680CD goat anti-mouse (red; Cat: 926-68070) at 1:20,000. The membrane was then imaged via a LICOR Odyssey scanner and quantified with Image Studio Lite software as “signal” (intensity with background subtracted). For full SOPs, please visit www.birtwistlelab.com/protocols. Each signal (pERK, pAKT, pEIF4E-BP1, and cyclin D) was first divided by the respective α-Tubulin signal (loading control) in the same lane to normalize for lane-to-lane differences in loading. Then to normalize for any systematic inter-well differences, each ratio was then normalized to the serum-starved control lane in its well. Raw images can be found in Figure S4.

##### BrdU incorporation/propidium iodide flow cytometry experiments and analysis

MCF10A cells were seeded at ∼1 million cells in a 10cm dish, and incubated overnight in full growth medium. The next morning, the full growth media was aspirated, cells were washed once with PBS, and then placed in starvation medium for 24 hours. Cells were treated with indicated doses of EGF (10nM), Insulin (10μg/mL), the combination dose (10nM EGF + 10μg/mL Insulin), and 30 min pre-incubation with a MEK inhibitor (PD0325901, Sigma, Cat: PZ0162) or AKT inhibitor (MK2206, ChemieTek, Cat: CT-MK2206) as indicated (both at 10μM). An hour before lifting, cells were treated with a pulse of 10 μM BrdU (Sigma). Cells were trypsinized, washed with 4°C 1X PBS and pelleted by centrifugation at 1000rpm for 5 minutes. To fix cells, they were resuspended in 200uL 4°C 1X PBS and 2mL of 4°C 70% ethanol (v/v in ddH_2_0) were added dropwise while vortexing at low speed. The cells were then stored at 4°C for at least 30 minutes. Cells were pelleted at 1000rpm for 5 minutes and resuspended in 1mL PBS. We added 1mL of 4N HCl and incubated the cells at room temperature for 15 minutes. We washed the cells by centrifuging at 1000rpm for 5 minutes and resuspending in 1mL PBS. After another spin, we resuspended the cells in 1mL PBT (PBS + 0.5% BSA (g/L) + 0.1% Tween 20 (v/v) (Sigma)). We spun the cells down and resuspended in 200 μL of PBT containing 1/40 dilution of anti-BrdU antibody (BD Biosciences) and incubated at room temperature for 30 minutes. We then added 1mL of PBT, spun the cells down and resuspended in 200uL PBT containing 1/40 dilution of anti-mouse FITC conjugate (BD Biosciences); we incubated cells in this for 30 minutes at room temperature in the dark. We added 1mL of PBT, spun cells down and resuspended in 250 μL PBS containing 250μg/mL RNAseA (Qiagen MaxiPrep Kit) and 10μg/mL propidium iodide (Sigma). We incubated the cells at room temperature for 30 minutes in the dark prior to analysis by flow cytometry on an Accuri C6 (BD Biosciences). We first gated for non-debris cells (FSC-A vs. SSC-A), then gated for singlets (SSC-A vs. SSC-H), then plotted PI intensity (x-axis) versus BrdU intensity (y-axis). We quantified percentages of cells in G1/G0 phase (First PI peak, BrdU negative), S-phase (BrdU positive), and G2/M phase (2^nd^ PI peak, BrdU negative). Our metric of percent cell cycling was the S-phase population plus the G2/M population percentages. Raw data and gates are available via Cytobank upon request.

##### Annexin V/Propidium Iodide flow cytometry experiments and analysis

MCF10A cells were seeded at ∼30% confluence in 12-well plates (75,000 cells / well), and incubated overnight in full growth medium. The full growth medium was replaced by starvation medium for 24 hours. Cells were then treated with different growth factor-inhibitor combinations as indicated and cells were assayed at 24, 48, and 72 hours post stimulation. A MEK inhibitor (PD0325901, Sigma, Cat: PZ0162) and/or AKT inhibitor (MK2206, ChemieTek, Cat: CT-MK2206) was used at 10μM. Anisomycin (Sigma, Cat: A9789), an inhibitor of protein and DNA synthesis, used at 10μM served as a positive control for cell death (death percentages were always greater than 90% cell death). Cells in full growth medium without treatment served as a negative control (death percentages were always less than 10% cell death). Cells were trypsinized, resuspended in PBS and pelleted by centrifugation at 1100g (to ensure collection of dead cells) for 5 minutes at room temperature. Cells were then resuspended in 100uL of Annexin-V binding buffer (10mM Hepes, 140mM NaCl, 2.5mM CaCl_2_ (Sigma) in ddH_2_0 pH 7.4) with 5uL of AnnexinV-FITC (BioLegend, Cat: 640906) and 0.1μg/mL propidium iodide (Sigma). Cells were incubated for 20 minutes at room temperature prior to analysis by flow cytometry. In TRAIL experiments, cells in full growth medium were treated with multiple doses of superkiller TRAIL (tumor necrosis factor (TNF)-related apoptosis-inducing ligand) (Enzo Life Sciences, Cat: ALZ-201-115-C010). After 5-hour TRAIL treatment, cells were harvested and analyzed by flow cytometry as described above, and a dose-response curve of two-fold serial dilutions of TRAIL ranging 2-300ng/mL was plotted. We first gated for non-debris cells, then gated for singlets (both as above), then quantified the % of cells in the Annexin-V positive population, which was our metric for percent cell death. Raw data and gates are available via cytobank upon request.

##### Etoposide flow cytometry experiments

MCF10A cells were seeded in full growth medium at approximately 30% confluence in 12-well plates (75,000 cells / well) and incubated overnight. Cells in full growth medium were then treated with 100 μM of Etoposide (Cayman Chemical, Cat: 12092) and assayed at 24, 48, and 72 hours post stimulation. Alternatively, after the cells attached overnight, the full growth medium was aspirated, cells were washed once with PBS, and then placed in starvation medium for 24 hours. These cells were then treated with 100 μM Etoposide and assayed at 24 and 48 hours post stimulation. Again, Anisomycin (10μM; Sigma, Cat: A9789) was used as a positive control for cell death and cells in full growth medium without treatment were used as a negative control for cell death. Cells were assayed with Annexin V-FITC and PI and analyzed as described above for the apoptosis flow cytometry experiments. Raw data and gates are available via cytobank upon request.

#### Computational Methods

##### Model Implementation

The model was implemented and simulated using MATLAB R2014b. We have noticed that later versions of MATLAB are not yet compatible with our model so using different versions may require additional programming. The model switches between an ordinary differential equation portion and stochastic gene expression portion with a 30 second time step. We used a MATLAB interface to CVODES (sundialsTB v.2.4.0), which greatly increased computational speed for the deterministically simulated model portion (ODEs). Most simulations were run on a MacBook Pro with a 2.5 GHz Intel Core i5 processor and 16GB of RAM. All code is available upon request or, upon publication downloadable from www.birtwistlelab.com.

##### Initial concentrations

In order to tailor the model to the expression context of MCF10A cells, we acquired gene copy number information from a literature source (Bessette et al., 2015), mRNA expression levels by performing RNA-seq, and protein expression levels by performing mass spectrometry-based proteomics. We acquired RNA-seq and proteomics data on serum-starved cells. We used proteomics data to set initial protein levels for the RTK, proliferation and growth, and apoptosis submodels. For the majority of proteins in the cell cycle and DNA damage submodels, we used initial conditions from their original source models. One reason is that the serum-starved condition (at 24 hours) is non-cycling and low stress, with little relevant expression of such proteins. Another reason is because these submodels already included intrinsic synthesis and degradation reactions, which we largely did not alter. As exceptions to this rule, we set initial cyclin D, p21, BRCA2, MSH6, and MGMT using proteomics data, as these were either proteins we added to the submodels *de novo* (all except cyclin D) or served as a link between submodels (cyclin D). We acquired a value for initial ribosome level from the literature; we found a range of values from ∼2 million to ∼10 million copies per cell across mammalian cells (Milo et al., 2009), thus we estimated 6 million copies of ribosomes copies per cell as an intermediate value. Note this value is ∼10-fold greater than the estimated number of mRNA molecules / cell, in line with the notion of the poly-ribosome (multiple ribosomes simultaneously translating the same transcript) and translation initiation being largely rate-limiting. The initial concentrations for all ligands and all post-translationally modified forms of proteins were initially set to zero. The levels of post-translationally modified forms of proteins were approximated during the initialization procedure (see Supplementary Methods for details).

##### Calculation of transcription and translation parameters

Each gene exhibits a unique transcription and translation rate, which we calculated from our RNA-seq and proteomics data. The general form of the deterministic differential equations describing transcription and translation are as follows:

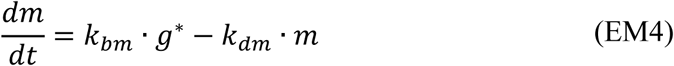

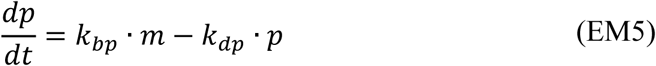

Where *g*\* is active gene levels, *m* is mRNA levels, *p* is protein levels, *k*_*bm*_ is the transcription rate constant, *k*_*bp*_ is the translation rate constant, *k*_*dm*_ is the mRNA degradation rate constant, and *k*_*dp*_ is the protein degradation rate constant. At steady state, the transcription and translation rate constants are given by

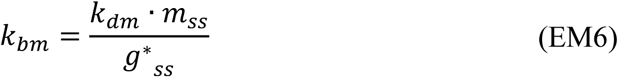

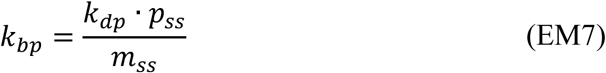

The mRNA and protein degradation rate constants were calculated from gene-specific half-life measurements observed in data from (Schwanhäusser et al., 2011a).

Measurements from Schwannhausser et al. were generated in NIH-3T3 cells, a murine fibroblast cell line; therefore, how do we know these mRNA and protein half-lives will apply to MCF10A human breast epithelial cells? First, observations point to consistent gene-specific mRNA-to-protein ratios across different human cell types (Wilhelm et al., 2014), which, we reason, supports the conservation of mRNA and protein half-lives across distinct cell types. Second, previous studies have reported conservation of mRNA and protein half-lives across species (Friedel et al., 2009; Schwanhäusser et al., 2011b). These reports give us confidence that mRNA and protein half-lives taken from Schwannhausser et al (2011) and other mammalian cell sources serve as reasonable estimates for half-life estimates in our MCF10A context.

We quantified the levels of mRNA and protein at steady-state (*m*_*ss*_ and *p*_*ss*_, respectively) based on our RNA-seq and proteomics measurements. The amount of active gene product at steady-state (*g*\* _*ss*_) was derived from equations. The differential equation for active gene production over time, given summation over a large population of cells, is defined as:

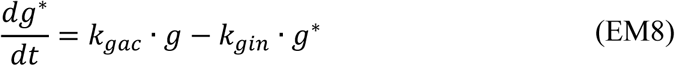

Applying moiety conservation to gene amount yields the following steady-state condition

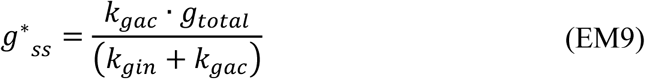

Gene activation and inactivation rate constants were obtained from the literature (Suter et al., 2011) (see Supplementary Methods for more details). As mentioned above, the total amount of gene copies for each gene (*g*_*total*_) was also obtained from a literature source (Bessette et al., 2015).

##### Creating a simulated cell population in silico

In an experimental in vitro setting, cells on a dish exhibit cell-to-cell variability in mRNA and protein expression levels. This natural variability is largely due to stochastic gene expression processes and can in some cases lead to cell-to-cell differences in phenotypic response to a perturbation. We desired to mimic this natural variability in our computational simulations. Because the model begins with the same initial conditions for each cell, we first run a stochastic simulation for each cell for 24 hours in the absence of external stimuli prior to performing any in silico experiment. This initialization step introduces a natural amount of variation in protein and mRNA levels across the simulated cell population, which randomizes the serum-starved condition in subsequent simulations (see Figure 2B).

##### Lasso regression and support vector machine (SVM) training

As part of our analyses we performed two lasso regressions; one related to apoptosis and one related to cell cycle. We used lasso regression to reduce our set of predictors for the SVM training. For apoptosis, we desired to identify genes that were predictive of whether a cell would die early (before 40 hours) or late (after 40 hours) in response to dual MEK and AKT inhibition. We used the `lasso` function in MATLAB to perform the regression. As predictor variables we used the total observable quantities of each protein (sum across all forms of a protein, i.e conserved moiety), either at the initial time point, or averaged across a certain time frame (8 or 40 hours). Our response variable binary. We used the default settings and set the “CV” parameter (fold cross validation) to 5. Time-averaged Bim and Bcl2 were the top two hits. We then trained a binary SVM classifier using the function `fitcsvm` in MATLAB, using only time-averaged Bim and Bcl2 total protein levels as predictor variables and early/late death responses as the response variables. We set the “Standardize” option to “true”, which normalizes the predictor data. We used a total of 200 cells for training and another 200 for testing. We tested the model using the `predict` function and generated ROC curves using the `perfcurve` function in MATLAB. For the cell cycle analysis, we performed the same steps as above though only considered the initial concentrations of each protein as predictors. Our response variable was binary depending on whether the cell divided within the 30-hour time course or not. CRaf and BRaf were the top two hits from the lasso regression, which we then used to train a binary SVM classifier, and generate ROC curves as above.

### QUANTIFICATION AND STATISTICAL ANALYSIS

#### Significance Testing

Significance testing was performed by using the `ttest2` function in Matlab. Three stars indicate a p-value less than 0.001, two stars indicate a p-value in-between 0.001 and 0.01, and one star indicates a p-value between 0.01 and 0.05. Error bars correspond to standard error of the mean where possible.

#### Flow Cytometry Software

Analysis of flow cytometry data was performed using Cytobank. Raw data and gates are available via cytobank upon request.

### DATA AND SOFTWARE AVAILABILITY

The MATLAB code used to run the model is available upon request or upon publication downloadable from www.birtwistlelab.com.

### ADDITIONAL RESOURCES

A full SOP for the μ-Western blot procedures can be found at www.birtwistlelab.com/protocols. More detailed SOPs for RNA isolation (DToxS SOP A – 1.0: Total RNA Isolation), RNA-seq library preparation (DToxS SOP A – 6.0: High-throughput mRNA Seq Library Construction for 3’ Digital Gene Expression (DGE)), RNA sequencing (DToxS SOP A – 7.0: Sequencing 3’-end Broad-Prepared mRNA Libraries), and analysis (DToxS SOP CO – 3.1: Generation of Transcript Read Counts) can be found at www.dtoxs.org.

## Chapter 1: Model Overview

### 1.1 Pathway and submodel selection

We sought to build a model could integrate the major cancer signaling pathways. A pancancer analysis by The Cancer Genome Atlas (TCGA) identified four major pathways that are frequently mutated across a variety of human cancers. These are the RTK-Ras-Raf pathway, the PI3K-AKT-mTOR pathway, cell cycle pathways, and p53-DNA repair pathways (Ciriello et al., 2013). To this list we added apoptosis pathways, in order to simulate cell death responses, and gene expression and degradation processes. With these pathways in mind, we created six “submodels”: (1) receptor tyrosine kinase (RTK) (Birtwistle, 2015; Birtwistle et al., 2007; Bouhaddou and Birtwistle, 2014; Kiselyov et al., 2009; Park et al., 2003), (2) proliferation and growth (which includes Ras-Raf-MAPK plus PI3K-AKT-mTOR pathways) (Birtwistle et al., 2007; von Kriegsheim et al., 2009; Nakakuki et al., 2010), (3) cell cycle (Gérard and Goldbeter, 2009), (4) apoptosis (Albeck et al., 2008), (5) DNA damage response (Batchelor et al., 2011), and (6) gene expression (the genes/proteins involved in each submodel are noted in main text Figure 1). To create our model, we assembled already-published models form the literature into a whole. The entirely of several submodels were already well represented by published models, such as cell cycle, apoptosis, and DNA damage response, to which we made very minor changes (details in Chapter 2). The RTK, proliferation and growth, and expression submodels were built largely from scratch, though prior models heavily inspired them (see above references). We considered models from the literature based on (*i*) whether they included proteins identified by TCGA as commonly mutated across cancers and (*ii*) whether they were built in conjunction with experimental observations in mammalian cells. All submodels were formatted according to the same rubric: lists of rate laws were extracted from each submodel and a stoichiometric matrix was created to define the elements of each differential equation. As a quality control measure, we ensured that we could reproduce the simulations from each original source model. Once each submodel was finalized, all reactions and stoichiometric matrices were assembled into an integrated system.

### 1.2 Model structural overview

The model is composed of 1197 total species, which includes all genes, mRNAs, lipids, proteins, and post-translationally modified proteins/protein complexes. There are a total of 141 genes that are considered across all submodels. Each gene possesses an inactive gene product, an active gene product, and an mRNA product (141+141+141=423 total species included in expression submodel). Because many of these genes are functionally redundant, these 141 mRNAs are summed during translation to create 102 “protein conglomerates”. Protein conglomerates represent functionally unique proteins (within the scope of our model) (see Supplementary Table 1 for list of genes and their mapping onto protein conglomerates). Once translated, these 102 protein conglomerates can become post-translationally modified in a number of ways; some can be phosphorylated, others form complexes, etc. This creates an additional 672 species across all submodels. These nascent proteins (102) plus their post-translationally modified versions (672) make up a total of 774 protein species (102+672=774). The 423 genes and mRNA species plus the 774 protein species comprise the 1197 total molecular products represented by the model.

We opted to represent the majority of reactions as elementary steps to the extent possible, using mass action kinetics. However, sometimes we employ Michaelis-Menten or Hill representations to capture events where the mechanisms are not sufficiently elucidated. Kinetic parameters are taken from original submodel sources, previous models, experimental studies, or estimated to fit empirical observations, many of which are discussed in the sections that follow.

### 1.3 Synthesis and degradation

The expression submodel controls the synthesis of most proteins in the model. The production of mRNA is based on a stochastic algorithm we devised (detailed in Chapter 2). Protein synthesis, 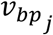, for each protein, *j*, is defined by (see Chapter 2 for derivation):

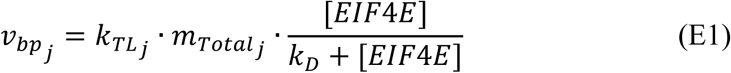

This equation is used to describe the synthesis of the majority of proteins in the model, with a few exceptions in the cell cycle and DNA damage submodels (details in Chapter 2).

#### Protein synthesis in cell cycle and DNA damage submodels

The original cell cycle (Gérard and Goldbeter, 2009) and DNA damage (Batchelor et al., 2011) models that we acquired already possessed their own synthesis reactions. Due to the dynamical complexity of these submodels, we needed a way to reconcile their original synthesis reactions with our equations. To do this, we required that, at steady state:

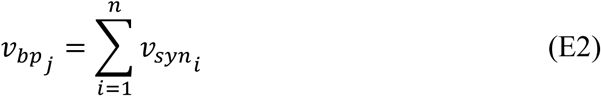

Where 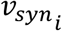 equals each original synthesis reaction, *i*, for a particular protein, *j*. For example, the synthesis of MDM2 is defined by a zero-order reaction plus a reaction that is dependent on the levels of active 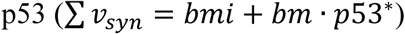. To meet this requirement, we defined 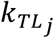, a translation rate constant for a particular protein, *j*, as a dynamic term that could account for changes in *v*_*syn*_. We thus define 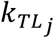 as:

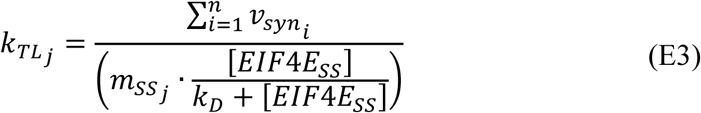

Here, 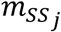 represents the levels of mRNA at steady state (from RNAseq data) for a given gene, *j* and *EIF*4*E*_*SS*_ are the levels of free EIF4E at steady state (defined during initialization procedure, described in Chapter 3). The final equation for protein production for species in the cell cycle and DNA damage submodels is:

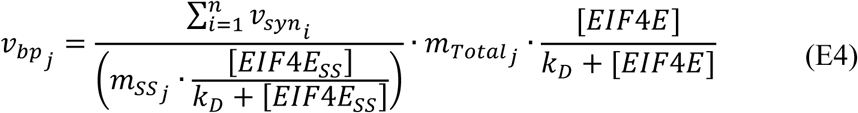

Where 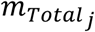 and *EIF*4*E* are levels of total mRNA for every gene, *j*, and free EIF4E currently in the system. By doing this, we allow for natural fluctuations in 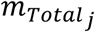 and *EIF*4*E* levels to affect the overall translation rate of proteins in the cell cycle and DNA damage submodel. However, as you will see (below and in Chapter 2.6) we did not allow the majority of cell cycle proteins to be stochastically regulated (with the exception of cyclin D and p21), as it resulted in non-biologically plausible behavior.

#### Lipid synthesis and degradation

Lipids represented in the model include phosphatidylinositol (3,4,5)-triphosphate (PI(3,4,5)P or PIP_3_) phosphatidylinositol 4,5-bisphosphate (PIP_2_), diacylglycerol (DAG), and inositol triphosphate (IP_3_). In our model, PIP_2_ is synthesized by a zero-order reaction. PIP_2_ can be cleaved by PLCγ to produce DAG and IP_3_. PIP_2_ can also be phosphorylated by PI3K to become PIP_3_. All lipids are degraded by first-order degradation reactions. Degradation rate constants for all lipids are taken from a literature source and references therein (Zhang et al., 2014). We increased the degradation of DAG by a factor of 10 to fit ERK signaling dynamics to experimental data.

#### Protein degradation

Proteins (protein conglomerates and post-translationally modified proteins) and transcripts follow first-order degradation kinetics. Protein and mRNA half-lives were taken from three sources, with preference given to the first source, then if a gene was not found to the second source, and so on: i) (Schwanhausser et al., 2011), ii) (Tani et al., 2012) (only for mRNA), iii) from miscellaneous literature references on mammalian cells (see Supplementary Table 1 for gene-specific half-life values and sources). Half-lives (τ) are converted into first-order degradation rate constants (*k*_*deg*_) using the following equation:

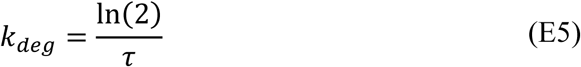

An additional layer of complexity arises when we consider the degradation of post-translationally modified protein monomers or protein complexes. In the absence of evidence for post-translational regulation of protein stability, degradation rates of post-translationally modified protein monomers were set equal to that of the non-post-translationally modified monomer. However, there are instances when, for example, the phosphorylated form of a protein may be degraded at a higher rate, due to the recruitment of degradation machinery, or, conversely, may be stabilized and thus degraded more slowly. For example, altered degradation rates have been shown to occur for cleaved caspases (Tawa et al., 2004), phosphorylated β-Catenin (Aberle et al., 1997), phosphorylated BIM (Luciano et al., 2003), phosphorylated BAD (Howie et al., 2008), and phosphorylated cFos and DUSP proteins (Nakakuki et al., 2010a and references therein), which we encoded in the model. Regarding the degradation of protein complexes, we assumed that the degradation rate of each complex is equal to the degradation rate of the most quickly degraded protein in the complex. Sometimes, however, the binding between two proteins can serve as a stabilizing force. We employ this reasoning to the degradation of phosphorylated cFos bound to cJun (AP1 complex), which we set equal to the literature-acquired degradation rate for phosphorylated cFos. We do this because phosphorylated cFos has been shown to be more stable compared to the relatively quick degradation rate for unphosphorylated cFos (Murphy et al., 2002; Nakakuki et al., 2010). Enabling AP1 to have an identical degradation rate also prolongs the AP1 signal in response to a mitogenic stimulus.

The cell cycle and DNA damage submodels possessed built-in degradation reactions, many of which had non-linear components. To retain the fidelity to the original model, we left degradation reactions in these submodels as they were. Exceptions to this rule include proteins we added to these models *de novo* or proteins that connect submodels; these are cyclin D, p21, BRCA2, MGMT, MSH6, and MDM4. The degradation reactions for these exceptions are modeled using the same first-order degradation reactions as in the other submodels.

#### Failure of incorporating stochastic gene expression into cell cycle submodel

During a stochastic simulation, all genes are simulated stochastically except for the majority of cell cycle genes. Cyclin D and p21 are the only two cell cycle genes that are simulated stochastically. Interestingly, incorporating stochastic gene expression processes into the entire cell cycle submodel created non-biologically plausible behavior. Principally, cells would randomly commit to cell division in the absence of a mitogenic stimulus. We attempted to alleviate this artifact through multiple modeling additions including increased availability and number of cyclin dependent kinase inhibitors, but could not find the solution. Therefore, to prevent this behavior, we only permitted stochastic control of cyclin D and p21 in our model; the other cell cycle proteins were simulated deterministically. Of course, stochastic variation in cyclin D and p21 levels propagate through to other cell cycle proteins, which introduces a much smaller degree of noise into the submodel compared to when each gene is simulated stochastically. We also did not allow any variations in the levels of EIF4E to affect cell cycle proteins other than cyclin D and p21. This result suggests that there may be hitherto unknown mechanisms by which the cell prevents against random cycling other than those formalized in this model.

### 1.4 Hybrid stochastic-deterministic algorithm

We desired to integrate stochastic gene expression processes into our model in order to capture natural fluctuations in protein levels that may render a cell more or less sensitive to a particular stimulus. We designed a hybrid stochastic-deterministic modeling framework in order to increase computational efficiency above the low efficiency expected of a fully stochastic algorithmic approach. Although the number of gene copies in a cell (roughly 2 copies) and the number of mRNAs in a cell (∼20 copies per cell on average) are relatively low, and thus amenable to efficient stochastic simulation, the number of protein molecules can approach hundreds of thousands of molecules per cell, rendering stochastic simulation of them inefficient. Conveniently, however, as molecule number approaches that typical of protein levels, the effects of stochasticity become small to negligible. Thus, we decided to simulate gene switching and mRNA birth-death stochastically and all protein processes (e.g. phosphorylation, dimerization, protein degradation, etc.) deterministically. Updated mRNA quantities were passed from the stochastic to the deterministic component. Signaling entities that act as transcriptional activators or repressors were transferred in the reverse direction, from the deterministic to the stochastic component. Stochastic implementation was based on Poisson processes (details in Chapter 2.1).

A critical concern was defining a time step between the stochastic and deterministic components that balanced computational efficiency with the need to capture the maximum number of stochastic gene switching events. We selected a time step of 30 seconds, which was the maximum time step ensuring (*i*) the probability of two gene switching events occurring within a single time step was low (below 0.1%; Figure S2A) and (*ii*) phenotypic behavior was not altered, such as timing of cell cycle entry (Figure S2B). This latter point was evaluated during model development.

### 1.5 Stoichiometric matrix formulation

In order to simplify and organize model implementation, we use the stoichiometric matrix formalism (Jeong et al., 2000; Schilling and Palsson, 1998). We created a matrix that describes the stoichiometry (either a 0, 1, -1, 2, or -2) between species (rows) and reactions (columns) in the model. The left hand side of the system of ODEs at each time step is calculated using the following matrix equation:

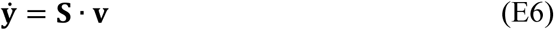

If *n* is the number of species and *m* is the number of reactions, then **S** is an n-by-m stoichiometric matrix, **v** is an m-by-1 vector of rate laws, and 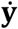 is an n-by-1 vector of the first time derivatives of model species concentrations. We modified this basic form to correct for cell compartment volumes (see Section 1.6, below, about Compartments).

### 1.6 Compartments

The model contains four compartments with associated volumes, *V*: extracellular space (*V*_*e*_), cytoplasm (*V*_*c*_), nucleus (*V*_*n*_), and mitochondria (*V*_*m*_). A typical mammalian cell has a volume between 100-10,000μm^3^, or 0.1-10pL (Milo et al., 2009). As a middle ground and based on visual inspection of phase contrast images, we approximated our total cell volume (*V*_*T*_) as 7pL (Ballesta et al., 2014; Geltmeier et al., 2015). Based on dimensions from phase contrast images, we determined the nucleus to be roughly a quarter of the cell’s volume (thus, *V*_*n*_ = *V*_*T*_/4). We then calculated the size of the cytoplasm to be equal to the difference between the total cell volume and nuclear volume (*V*_*c*_ = *V*_*T*_ - *V*_*n*_). The mitochondrial volume was previously estimated to be approximately 7% of the size of the cytoplasm (Albeck et al., 2008). For cell culture experiments, the volume of the extracellular space is much larger than the volume of single cells. We set the volume of the extracellular space to be 50μL, which is the volume of media typically placed into one well of a 96-well plate. Although experiments are sometimes done in different vessel sizes, this large volume ratio renders simulations insensitive to typical variation across experimental culture conditions.

Based on their cellular localization, every species and every reaction was assigned one of these four home compartments. Importantly, when modeling in terms of concentration, one must introduce volume corrections when (*i*) reactants in a rate law possess different home compartments or (*ii*) rate laws comprising a differential equation possess different home compartments. When a reaction possesses reactants with different home compartments, we introduce a volume correction term to volume correct the outsider reactant to be defined in terms of the home compartment of the reaction, as such:

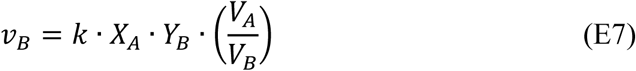

Where *X*_*A*_ is a species in a compartment of volume *V*_*A*_, *X*_*B*_ is a species in a compartment of volume *V*_*B*_, and *v*_*B*_ is a reaction with a home compartment of volume *V*_*B*_. For example, if we consider the reaction describing the binding of EGF to EGFR, which takes place in the extracellular space, the home compartment of EGF is the extracellular space (*V*_*e*_) whereas the home compartment of EGFR is the cytoplasm (*V*_*c*_). In this case, we correct the concentration of EGFR to be in terms of the extracellular volume by multiplying the first order reaction by *V*_*c*_/*V*_*e*_.

In addition to this bimolecular volume correction on the reaction level, sometimes an ordinary differential equation can possess reaction terms that are of different home compartments. To correct for this, we multiply every reaction by its respective home compartment to remove units of volume. We multiply this vector of “volume-less” reaction values, **v**_**VC**_, by our stoichiometric matrix, producing an n-by-1 vector 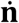 with units of nmol / time (and length equal to the total number of species, *n*):

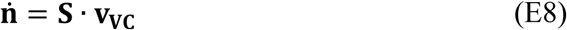

Finally, to add units of volume back to our 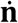 quantities, we element-wise divide by the home compartment of each species, which produces our final n-by-1 vector of the first time derivatives of model species concentrations, 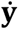.

### 1.7 Ribosomes and cell size

A cell must approximately double its contents prior to cell division; else, repeated rounds of cell division would eventually push a cell population to non-existence. One predominant mechanism for this is an increase in the number of ribosomes, which increases the translation rate globally across all proteins. We thus use ribosome quantity (related to ribosome number and not concentration) as a proxy for cell size. This has the effect of keeping ribosome concentration constant, which we lump into the effective translation rate constant. Therefore we are not able yet to account for how stochastic fluctuations in ribosome levels affect protein expression noise. However, since the average number of ribosomes in a mammalian cell, at ∼6,000,000 molecules per cell (Milo et al., 2009), is much greater than the average number of mRNA molecules, at ∼400,000 molecules per cell (Alberts, B. Johnson, A. Lewis, J. Raff, M. Roberts, K. Walter, 2008), we expect such fluctuations to be small compared to stochastic gene switching and mRNA birth-death, which we account for.

The de novo synthesis of ribosomes for cell growth and proliferation has been linked to activation of p70 S6K (Chauvin et al., 2014). This, on top of synthesis (zero-order) and turnover (first-order) terms for ribosome homeostasis, gives rise to the following differential equation for ribosome dynamics:

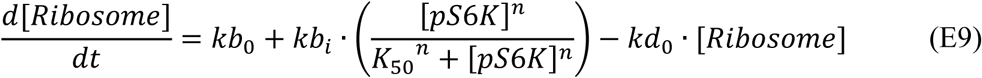

We determined the degradation rate for ribosomes by taking the average of the half-lives reported by Schwanhausser et al (2011) for all 40S and 60S ribosomal proteins.

To calculate *kb*_*i*_, the rate constant governing the upregulation of ribosomes by *pS*6*K*, we assumed that *pS*6*K* levels in the serum-starved state are low (approximately ∼1% of the total pool of S6K). In a highly mitogenic environment, we assume S6K will be highly phosphorylated (∼50%). We then estimated the value of *kb*_*i*_ (value of 0.04 nM^-1^s^-1^) such that, in response to a mitogenic stimulus, ribosome numbers would double during the course of one cell cycle (Figure S2D), which for MCF10A cells was determined to be between 16 and 24 hours (Albeck et al., 2013).

## Chapter 2: Submodel Methods

Here we describe more details about each submodel. Therein, we detail any changes we made to the original source models, where applicable. We also discuss the major biological behaviors we desired that each submodel possess. These are outlined as follows:

| Submodel | Required Properties |
| --- | --- |
| Expression | • Model is tailored to genomic, transcriptomic, and proteomic context of MCF10A cells.<br>• Stochastic gene expression is simulated with a computationally efficient algorithm.<br>• Cell-to-cell variability in mRNA and protein levels matches experimental observations.<br>• EIF4E levels possess extrinsic control over the translation rate.<br>• Ribosomes double during the course of one cell cycle. |
| Receptor Tyrosine Kinase | • Ligand-receptor cooperativity matches experimental observations<br>• Receptor trafficking kinetics reflects experimental observations. |
| Proliferation & Growth | • Receptors possess unique pathway preferences that match experimental observations.<br>• Basal, serum-starved state activity flux through ERK and AKT pathways exists.<br>• Dynamic dose responses of ERK, AKT, and mTOR signaling matches experimental data. |
| DNA Damage | • Original delayed differential equations are converted into ordinary differential equations.<br>• p53 dynamics corresponding to single- and double-stranded DNA breaks matches experimental observations.<br>• Rate of DNA damage repair is dependent on levels of repair enzymes.<br>• p53 activation dynamics exhibit “digital” and not “analog” behavior, whereby the number of p53 pulses, but not pulse height or width, scales to magnitude of DNA damage.<br>• Etoposide-induced DNA damage is dependent on the cell cycle stage (S-phase). |
| Apoptosis | • Robustness against small death signals.<br>• Model exhibits all-or-nothing death response when apoptosis signaling surpasses threshold. |
|  | • Dose and dynamics of TRAIL-induced extrinsic apoptosis matches experimental observations.<br>• Intrinsic apoptosis signaling responds to interrupted survival signaling and DNA damage induced upregulation of pro-apoptotic proteins. |
| Cell Cycle | • Cell cycle entry is driven by induction of cyclin D mRNA.<br>• Order and timing of cyclin/cdk complexes matches established observations.<br>• Cell cycle duration matches duration measured in MCF10A cells.<br>• Upregulation of p21 (a cyclin dependent kinase inhibitor) arrests the cell cycle. |

### 2.1 Expression

#### About

The expression submodel describes genes switching between an inactive and active state, transcription of the active gene to create mRNA, and translation of the mRNA to create protein. The mRNA and protein products can be degraded. Some genes in the model possess explicitly coded transcriptional activators and/or repressors, which can affect the transcription rate (see equations below). The translation rate of an mRNA is dependent on the mRNA concentration in the cell as well as on the levels of EIF4E, a rate-limiting cap-dependent translation initiation factor. Importantly, each mRNA and protein possesses a unique synthesis and degradation rate, resulting in dynamical behaviors that may play important roles in constraining protein function and phenotypic outcomes. Moreover, these gene-specific attributes are known to vary significantly across the genome (Schwanhausser et al., 2011). Therefore we sought to capture these gene-specific attributes to the extent possible.

Transcription and translation rates are estimated as part of the initialization procedure (see Chapter 3). mRNA and protein degradation rates are calculated from half-life measurements found in the literature (Supplementary Table 1). Although these half-lives were procured from different mammalian cell lines and cell types, we assume mRNA and protein half-lives to be fairly consistent between cell types. This is inspired by (*i*) the fact that gene-specific ratios of mRNA-to-protein are fairly consistent between different human cell types (Wilhelm et al., 2014) and (*ii*) evidence from several studies that report conservation of mRNA and protein half-lives across species (Friedel et al., 2009; Schwanhausser et al., 2011). These points give us confidence that mRNA and protein half-lives taken from other mammalian cell contexts serve as reasonable approximations of half-life estimates for MCF10A cells, and that altering protein levels through translational means, as we implement, is preferred. Finally, this submodel possesses a stochastic component; specifically, in gene switching and mRNA births and deaths. Translation and protein degradation are simulated deterministically, though fluctuations from gene expression propagate into protein levels over time. As described above, the model switches from the stochastic to the deterministic component (and back again) every 30 seconds of simulation time.

#### Epigenetics

Rates of gene switching were acquired from an experimental study (Suter et al., 2011). The study established a transgenic cell line expressing a short-lived luciferase from an unstable mRNA, allowing them to measure gene activation and inactivation kinetics for a handful of genes. The paper defined an average gene activation and inactivation rate, which we used for each gene in our model, similar to other modeling work (Bertaux et al., 2014) (see Table SI1).

**Table SI1.** Rate constants for gene activation and inactivation.

|  |  |
| --- | --- |
| $k_{gac}$ (gene activation) | $0.03 \text{ min}^{-1}$ |
| $k_{gin}$ (gene inactivation) | $0.3 \text{ min}^{-1}$ |

We model gene switching as a Poisson process. The Poisson distribution is a discrete probability distribution that describes the probability that a given number of events will occur within a certain time interval, independently from when the last event occurred. It is parameterized by lambda, which is defined as the expected number of events in the considered time interval. For gene switching, lambda is defined as the rate of gene activation, or inactivation, multiplied by the time interval:

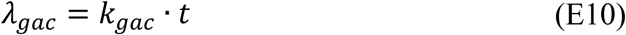

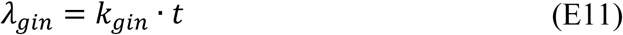

For each gene, we calculate the probability of getting zero switching events using the `poisspdf` function in Matlab evaluated at zero. This defines two probabilities: p_off_ (probability that a gene that was on would stay on; calculated using λ_*gin*_) and p_on_ (probability that a gene that was off would stay off; calculated using λ_*gac*_). Next, a random number was generated for each gene between 0 and 1 using the `rand` function. For genes that were on, if their random number was greater than or equal to p_off_, then they were switched from on to off. Similarly, for genes that were off, if their random number was greater than or equal to p_on_, then they were switched from off to on.

#### Transcription

Transcription and mRNA degradation were also modeled as Poisson processes. The number of mRNA births and deaths per time step were randomly sampled from Poisson distributions using the `poissrnd` function in MATLAB. Again, lambda terms were defined as the transcription rate or degradation rate of a gene multiplied by the time interval. The rate of transcription was determined by summing a leak term, which accounts for all non-modeled or constitutive transcription, and an induction term, which accounts for all modeled transcriptional induction. We define the rate law for transcription as:

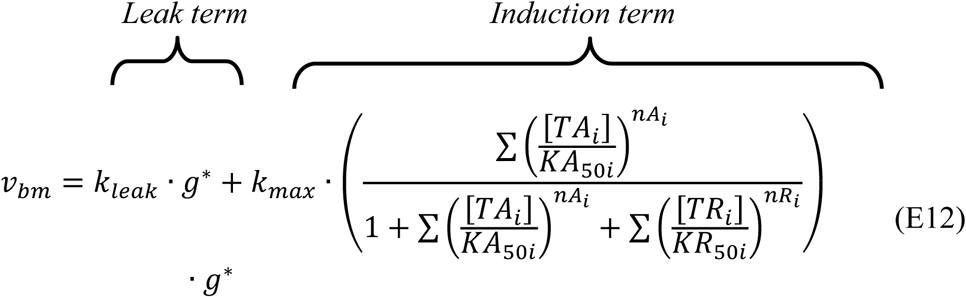

Here, *k*_*max*_ is the maximal transcription rate of RNA polymerase (same for all genes), which we determined to be 0.1 molecules per second (Iyer and Struhl, 1996; Kugel and Goodrich, 2000). For every transcriptional activator (*TA*_*i*_) or repressor (*TR*_*i*_), *i*, *KA*_50*i*_ and *KR*_50*i*_ are the concentrations needed to achieve their half-maximal effect on the transcriptional output, respectively. *nA*_*i*_ and *nR*_*i*_ values are hill coefficients for activators and repressors, respectively. K_50_ and hill coefficient values were tuned as part of model training. Hill functions such as these are commonly used to describe transcriptional induction terms (Alon, 2007; Nakakuki et al., 2010; Thattai and van Oudenaarden, 2001). The number of active or inactive genes in the cell are denoted by *g*\* or *g*, respectively.

The rate law for mRNA degradation is a first order term:

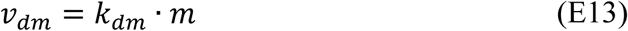

Here, *k*_*dm*_ is a rate constant derived from mRNA half-life data taken from the literature for each mRNA (Supplementary Table 1), and *m* is the number of mRNA molecules in the cell. Thus, the lambda terms for the Poisson distributions defining transcription and mRNA degradation are:

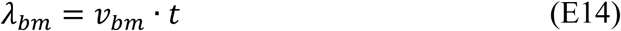

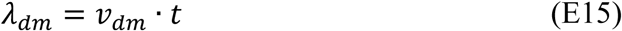

#### Translation

Equations describing translation are simulated deterministically. We derived a differential equation for translation that accounted for extrinsic control based on the levels of EIF4E in the cell, a critical initiation factor that binds to the mRNA 5’ cap to recruit it to an available ribosome (Mamane et al., 2004). Because translation initiation is thought to be the rate-limiting factor in protein production (Kudla et al., 2009; Sonenberg and Hinnebusch, 2009), we take the rate of translation initiation, and thus the rate of translation, to be linearly proportional to the amount of mRNA bound to an EIF4E molecule (*m*\*):

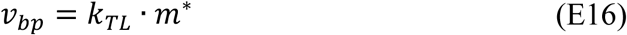

Here, *k*_*TL*_ implicitly includes the dependence on ribosome concentration, which is constant in our model because cell volume is proportional to ribosome number.

Next we sought to solve *m*\* in terms of the total amount of mRNA per cell, a metric that is experimentally quantifiable from the output of mRNA sequencing experiments. The differential equation describing the binding between free mRNA (*m*_*free*_) and EIF4E is defined as:

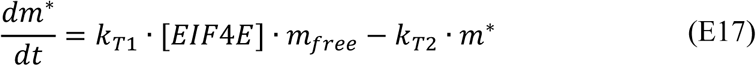

Solving for *m*\* at steady state yields:

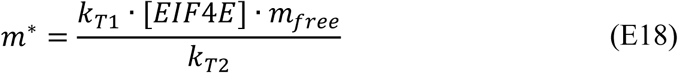

Given mRNA moiety conservation (*m*_*Total*_ = *m*_*free*_ + *m*\*), and the definition of the dissociation constant *K*_*D*_, yields

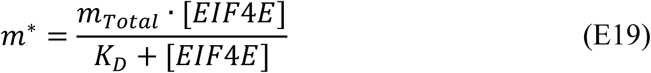

Plugging this definition for *m*\* back into Equation E16 gives our final equation for the rate of translation:

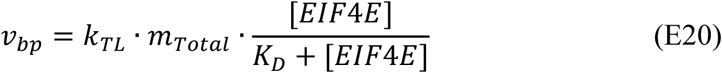

The final differential equation for proteins is also noted above in Equation E1.

### 2.2 Receptor Tyrosine Kinase (RTK)

#### About

The receptor tyrosine kinase submodel contains many receptors that are frequently overexpressed or mutated in cancer; these include the ErbB family (Her1-4), the hepatocyte growth factor receptor (cMet), the platelet-derived growth factor receptor (PDGFR), the fibroblast growth factor receptor (FGFR), the insulin-like growth factor receptor (IGFR), and the insulin receptor (INSR). For all receptors except for PDGFR, we allow receptors to form dimers in the absence of ligand, though at very low affinities. IGFR and INSR are exceptions, forming covalent cysteine bonds between receptor dimers prior to ligand binding and activation (Kiselyov et al., 2009).

Ligand stimulation induces receptor dimerization. Ligand can bind to either a receptor monomer or a preformed receptor dimer. Once a ligand binds a receptor monomer, it can dimerize with another receptor monomer or another ligand-bound receptor (forming either LRR or LRLR type species). The ligand binding to the preformed dimer can subsequently bind another ligand. Once receptor dimers are in complex with either one or two ligands, we term them signaling competent dimers (SCDs). For the ErbB family of receptors, EGF can only bind to EGFR, ErbB2 has no known ligand, and ErbB3 and ErbB4 can bind to heregulin. All family members can dimerize with one another, creating a plethora of possible combinations.

The formation of these SCDs induces the activation of the tyrosine kinase domain and the phosphorylation of intracellular tyrosine residues. This can recruit adaptor proteins, which mediates downstream signaling. For IGFR and INSR receptors, we require an additional step—binding to IRS—prior to phosphorylation. To construct the schematic for PDGFR, we used a simplified scheme presented by Haugh and colleagues (Park et al., 2003). In this scheme, the binding of ligand to PDGFR induces it to bind to another ligand-bound receptor. From this complex, one ligand can dissociate, which provokes the subsequent release of one receptor molecule. For a detailed model schematic please refer to Figure S1. All rate constant values, and sources for each rate constant, for the receptor tyrosine kinase submodel (as well as for the proliferation and growth submodel) can be found in Supplementary Table 2.

#### Ligand-receptor cooperativity

In order to reproduce appropriate dose-response behavior for each ligand-receptor system we ensured that the cooperativity behavior for each ligand-receptor system matched experimental observations from the literature (see table below). Specifically, we ensured the appropriate fold change between the K_D_ of the 1^st^ and 2^nd^ ligand-binding sites (see Table SI2). For example, it is thought that the binding of the first EGF molecule to EGFR occurs with ∼10 fold higher affinity than the binding of the second ligand (Alvarado et al., 2010). Because receptor cooperativity measurements can vary between cell types, we sought only that cooperativity behavior was in the correct category – either displaying negative, positive, or no cooperativity (Figure 3A and S3A in main text). As a general rule, we attempted to keep ligand-receptor association rate constants consistent across reactions and slower than diffusion encounter rate (we take them as 0.001 nM^-1^s^-1^). Ligand dissociation rate constants were tailored to match experimental observations as described. Also, wherever possible, we used generic receptor dimerization (0.01 nM^-1^s^-1^) and dissociation (0.001 s^-1^) rate constants, which serve as reasonable approximations for these reactions for which direct experimental measurement is difficult (Birtwistle, 2015). It was not always possible to match cooperativity and dynamics while satisfying detailed balance; this may be due to unmodeled but implicit energy dependent steps such as cycles of phosphorylation coupled to dimerization, or other receptor microdomain compartmentalization that is also energy dependent.

**Table SI2.** Receptor cooperativity parameters and literature sources.

| Ligand | Receptor(s) | Fold difference between binding sites | Cooperativity | Literature Sources |
| --- | --- | --- | --- | --- |
| EGF | EGFR | 10 | Negative | (Alvarado et al., 2010) |
| HRG | ErbB3, ErbB4 | 10 | Negative | (Hiroshima et al., 2012) |
| HGF | c-Met | 1 | None | (Takahashi et al., 1996) |
| PDGF | PDGFRα, PDGFRβ | NA * | Positive | (Park et al., 2003) |
| FGF | FGFR1, FGFR2 | 500 | Negative | (Mohammadi et al., 2005; Story et al., 1994) |
| IGF | IGF1R | 100 | Negative | (Christoffersen et al., 1994; |
|  |  |  |  | Kiselyov et al., 2009; De Meyts and Whittaker, 2002) |
| INS | INSR | 100 | Negative | (Christoffersen et al., 1994; Kiselyov et al., 2009) |

\* PDGFR was coded according to a simplified scheme depicted in a literature source (Park et al., 2003).

#### Receptor trafficking dynamics

Once receptors bind their cognate ligands and form dimers, they phosphorylate one another. Phosphorylated signaling competent receptor dimers can also be internalized and targeted for degradation, or can be recycled back to the plasma membrane from early endosomes. Receptor internalization and degradation is an important mechanism by which pathways shut off after acute stimulation (in addition to any negative feedback that might exist). Importantly, internalization and degradation kinetics can vary between receptor dimer types (see Table SI3). Of note, dimers containing EGFR are known to internalize and degrade 10X faster, and recycle back to the plasma membrane 10X slower, than when an EGFR molecule is not involved (Resat et al., 2003). Rate constants for internalization, degradation, and recycling were set to the following:

**Table SI3.**
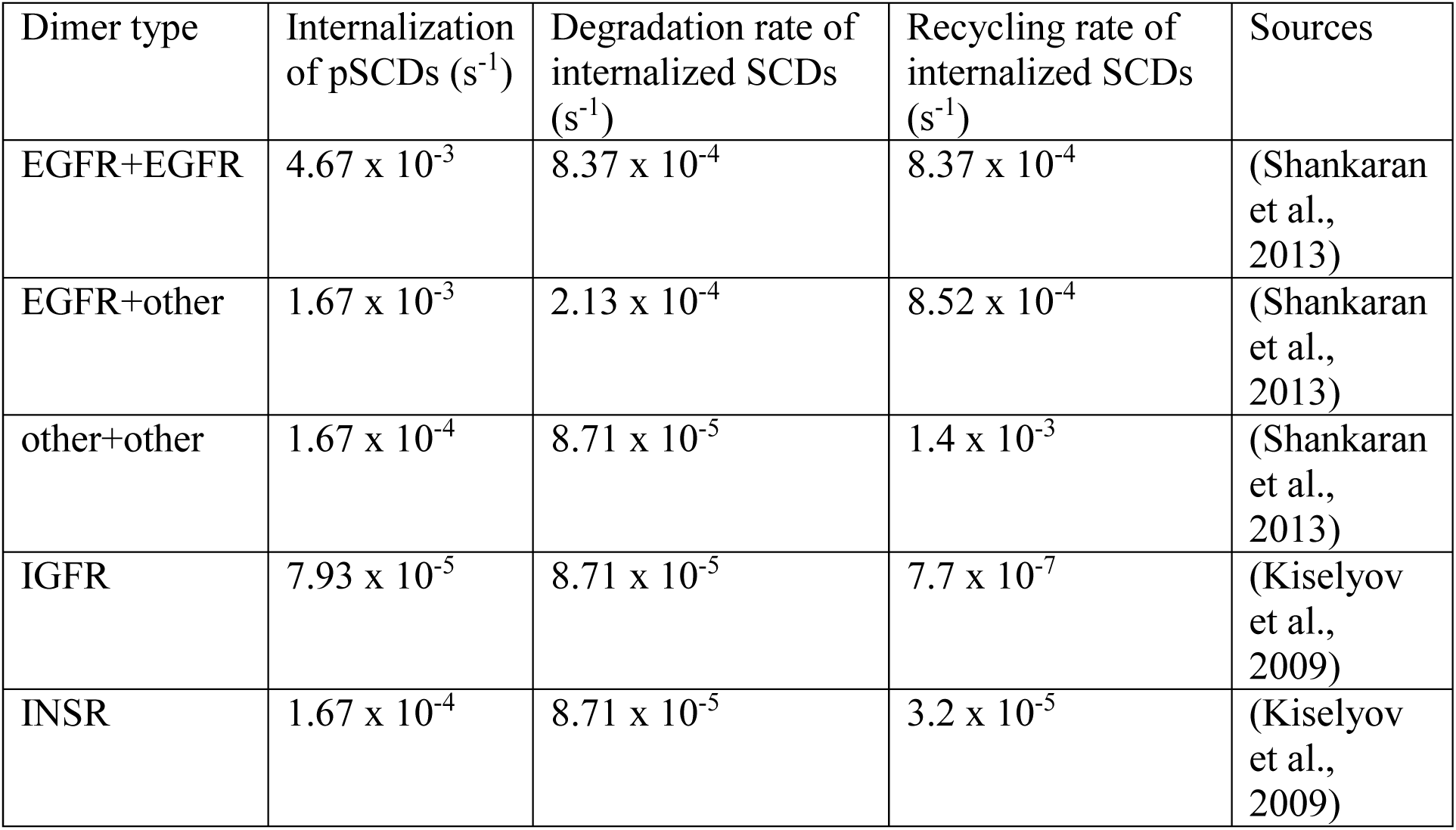
Receptor internalization and degradation kinetics.

It is thought that the early stages of endocytosis are clathrin-mediated; however, due to the limited availability of clathrin, this mechanism quickly becomes saturated. Once saturated, slower endocytosis mechanisms take over. To describe this process, our rate law describing receptor internalization contains two parts:

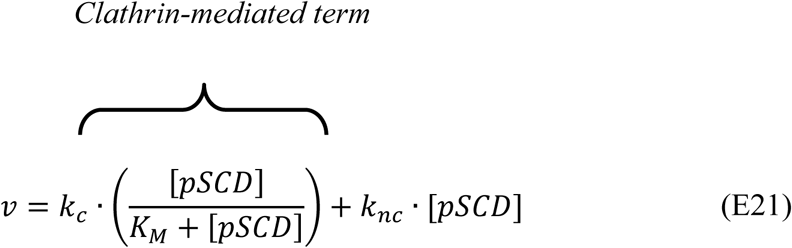

The first is the saturable, clathrin-mediated component, whose rate constant *k*_*c*_ indicates values described in Table SI3, above. The second part is a first order reaction whose rate constant *k*_*nc*_ is set to the “other+other” internalization rate for all receptor dimers. The term *pSCD* denotes the concentration of the phosphorylated signaling competent dimer and *K*_*M*_ is an effective Michaelis constant.

### 2.3 Proliferation and growth submodel

#### About

This submodel was largely build from scratch, with much inspiration from prior models (Birtwistle et al., 2007; von Kriegsheim et al., 2009; Nakakuki et al., 2010). The model depicts the binding of adaptor proteins to phosphorylated receptor tyrosine kinases, leading to the activation of two major signaling pathways—the ERK and AKT pathway. Once phosphorylated, receptors can become internalized and are known to signal to the ERK pathway from early endosomes (Burke et al., 2001; Gregory and Hann, 2000; McKay and Morrison, 2007). Many receptors then enter into late endosomes and are degraded; others are de-phosphorylated and recycled back to the plasma membrane (Resat et al., 2003). The activation of ERK (resulting in a cascade from Ras to Raf to MEK to ERK) results in the upregulation, phosphorylation and stabilization of cFos (with p90RSK), promoting it to bind cJun, forming AP1; active ERK also mediates the transcriptional upregulation of cFos (Murphy et al., 2002), dual-specificity phosphatases (DUSPs) (Brondello et al., 1997), and Sprouty proteins (Ozaki et al., 2001). There exists a positive feedback at the level of cJun, as AP1 (phosphorylated cFos bound to cJun) is known to transcriptionally upregulate cJun (Angel et al., 1988). In addition to the negative feedback ERK induces via DUSP phosphatases and Sprouty proteins, other mechanisms of negative feedback include ERK regulation of CRaf (Dougherty et al., 2005), receptor tyrosine kinases (Li et al., 2008), and Grb2-SOS (Chen et al., 1996) via direct (and/or indirect) phosphorylation.

The activation of AKT can induce the nuclear localization of β-Catenin through GSK3β, which is known to upregulate cMyc (He et al., 1998). Active AKT (along with ERK) can also activate mTOR signaling via the TSC complex. TSC complex inhibition by AKT leads to the upregulation of mTORC1, which is known to increase translation globally and regulate cell growth. This is accomplished by mTORC1-mediated phosphorylation and inhibition of EIF4E-BP1 (Manning and Cantley, 2003), causing it to dissociate from and make available EIF4E, as well as its phosphorylation and activation of p70S6K (Fingar et al., 2002), which, when active, drives ribosome synthesis. mTORC1 is also thought to exhibit negative feedback on PI3K (Carracedo and Pandolfi, 2008). As was done for the RTK reactions, here we also assume association reactions to be slower than diffusion limited (0.001 nM^-1^s^-1^). When tuning was needed (to match simulations to experimental observations), we principally altered dissociation rate constants and catalytic constants. As a baseline assumption, we set dissociation rate constants to 0.1 s^-1^ and catalytic constants for kinase/phosphatase reactions to 1 s^-1^ (Birtwistle, 2015). All rate constant values, and sources for each rate constant, for the proliferation and growth submodel (as well as for the RTK submodel) can be found in Supplementary Table 2.

#### Pathway preferences

Once intracellular tyrosine residues of signaling-competent dimers become phosphorylated, adaptor proteins are recruited, each of which leads to the activation of distinct downstream signaling pathways. The extent to which a receptor activates one pathway over another depends, in part, on the number of binding sites it possesses for adaptor proteins. In our model we consider three different adaptor proteins: Grb2 (and implicitly Shc), PLC-γ, and PI3K (composed of regulatory and catalytic subunits). We determined how many binding sites for each of these three adaptor proteins existed in each receptor based on known domains within each receptor sequence. To do this, we used ScanSite (http://scansite3.mit.edu) set to “medium” stringency setting (Obenauer et al., 2003). For Grb2 we summed across domains corresponding to Grb2 and Shc. For PI3K we summed across domains corresponding to PI3K SH2 and SH3 domains. For PDGFR we took the average between the number of sites on PDGFR-A and PDGFR-B isoforms for each adaptor. For FGFR, we averaged across FGFR1 and FGFR2 isoforms. For IRS, we averaged across IRS1 and IRS2 isoforms. These factors, depicted in Table SI4, served as multipliers in the rate laws describing the binding of phosphorylated signaling competent dimers to adaptor proteins.

**Table SI4.**
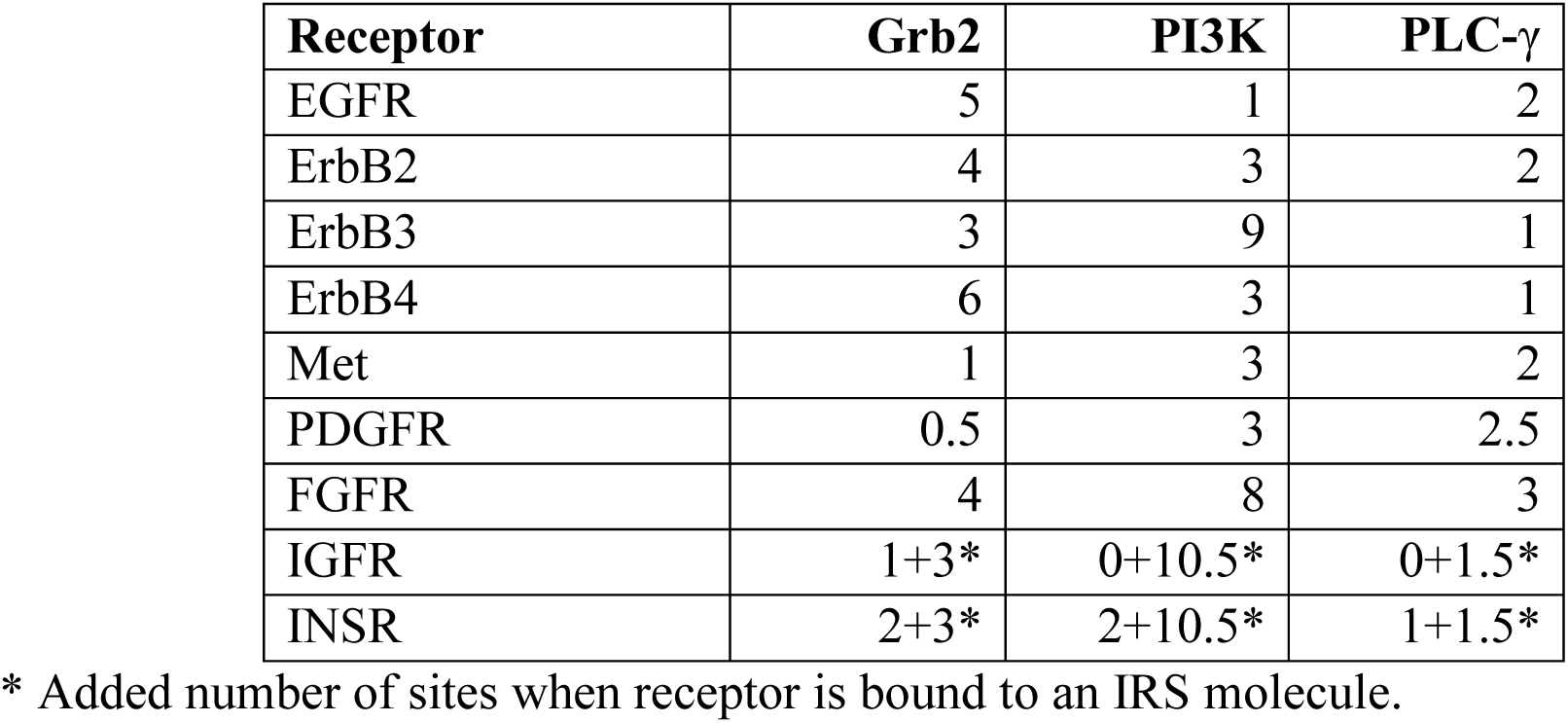
Number of adaptor binding sites on each receptor tyrosine kinase.

In addition, the affinity of adaptor proteins for phosphorylated tyrosine residues (Table SI5) was taken from a literature source that previously compiled data from multiple experimental studies (Birtwistle, 2015):

**Table SI5.** Affinity (K_D_) of adaptor proteins for phosphorylated tyrosine residues on receptors.

| | Grb2 | PI3K | PLC-γ |
| --- | --- | --- | --- |
| $K_D$ | 2600nM | 70nM | 510nM |

#### Basal fluxes

Even when cells are serum-starved, basal levels of signaling exist. We encode this basal flux by modeling basal activity of Ras (first-order Ras-GDP to Ras-GTP, and vice versa) and PI-3K (first-order PIP_2_ to PIP_3_, and vice versa). These propagate basal signals through the ERK and AKT (and mTOR) pathways. Normally these signals are between 1-10% of the total activation amplitude.

#### Dose and dynamic pathway responses

We stimulated MCF10A cells with various doses and combinations of EGF and insulin and measured pERK, pAKT, pEIF4E-BP1 (mTORC1 proxy) and cyclin D at various time points post-stimulation via μ-Western blot. We focused on EGF and insulin because we observed a synergistic interaction between the two growth factors and desired to investigate further (see Figure 6 in main text). Also, these two growth factors are critical for the growth of MCF10A cells. We tuned model parameters to match the dose and dynamic response of measured pERK, pAKT, and pEIF4E-BP1 levels at 0, 5, 30, 180, and 360 minutes post-stimulation (see Figure 3B-D in main text). In addition, we ensured that the proportions of simulated cyclin D levels at 6 hours post-stimulation for EGF (10nM), insulin (10μg/mL), and EGF plus insulin matched those measured via μ-Western blot (see Figure 6C in main text).

#### Incorporating inhibitors

Because many of our research questions revolved around how ERK and AKT activity related to various cell fate outcomes, several of our experimental perturbations included the inhibition of MEK (just upstream of ERK) and AKT. Our AKT inhibitor, MK2206, is an non-ATP competitive allosteric inhibitor, thought to prevent binding of the kinases that phosphorylate and activate AKT (Cherrin et al., 2010). Thus, we decided to model it binding to the non-phosphorylated form of AKT and inhibiting its activation. Conversely, the MEK inhibitor we use, PD-0325901, is thought to bind to the ATP-binding domain of MEK, preventing its ability to phosphorylate its targets (Barrett et al., 2008). We decided to code this as the MEK inhibitor binding to the doubly phosphorylated form of MEK.

### 2.4 DNA damage submodel

#### About

The DNA damage submodel describes the activation of ATR and ATM by single-and double-stranded DNA breaks, respectively. Single-strand breaks also at this juncture encapsulate point mutations that must be excised and replaced with the correct base pair by repair enzymes (although future models could take a finer grained approach). The activation of ATR and ATM leads to the production of sustained versus oscillatory p53 dynamics, as is known to occur in mammalian cells (Batchelor et al., 2011). The activation of p53 leads to the production of MDM2 and WIP1, both of which serve as negative feedback regulators on p53 activation—MDM2 is an E3 ubiquitin ligase that induces degradation of active p53 (Haupt et al., 1997); WIP1 has a negative feedback on ATM (Shreeram et al., 2006). DNA repair enzymes can repair DNA damage; here, we encode activities of BRCA2 (double-stranded breaks), MSH6 (single-stranded breaks), and MGMT (single-stranded breaks). p53 can also bind to, and be sequestered, by MDM4 (Bessette et al., 2015).

#### Converting DDEs to ODEs

The original DNA damage model from the Lahav lab (Batchelor et al., 2011) originally contained delay differential equations, which served to model a delay between p53 activation and the upregulation of WIP1 and MDM2. We replaced the delay differential equations with a series of 20 first-order ordinary differential equations that are meant to represent the mechanistic steps that are implied by the original delay (e.g. p53 biding to DNA, recruiting co-transcription factors, synthesizing RNA, etc.). These equations were of the form:

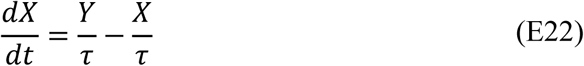

Here, Y would be the next variable in the cascade. We estimated the τ parameters using a genetic algorithm, implemented using the `ga` function in Matlab, which gave a near-perfect fit between the original model and ours (Figure SI1, below)

#### DNA damage induction by Etoposide

We also altered the way in which DNA damage was induced in the model. The original Bachelor et al. (2011b) model possessed an on or off switch for DNA damage. We decided to encode DNA damage as a continuous variable, one for single stranded breaks (SS) and one for double stranded breaks (DS). Here, we decided to focus on Etoposide as the principal DNA damage stimulus, a clinically approved inhibitor of topoisomerase II. Topoisomerase II is an enzyme that aids in the uncoiling of DNA during replication. Importantly, Etoposide is only lethal to cells actively traversing through S-phase (see Figure 4 in main text). We used the following equation to capture these properties:

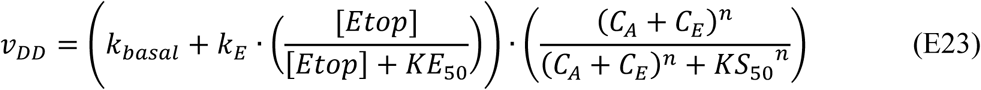

Here, *v*_*DD*_ is the rate of DNA damage accumulation (with equal contributions to the creation of single-and double-stranded breaks), *k*_*basal*_ is the basal DNA damage rate meant to reflect damage in the absence of an overt stimulus (estimated during initialization, see Chapter 3.4), *k*_*E*_ is the maximal rate of Etoposide-caused DNA damage for which *KE*_50_ is the half-maximal (estimated to fit model to cellular measurements), and *C*_*A*_ and *C*_*E*_ are the levels of active cyclin A/Cdk2 and cyclin E/Cdk2 in the model (these are the cyclins upregulated during S-phase and bound to their cognate cyclin dependent kinases).

#### DNA damage repair

For the repair of DNA damage, we incorporated DNA repair enzymes that could repair DNA in a concentration-dependent manner. The incorporation of DNA repair enzymes is important because they are frequently mutated in cancer. These genes included BRCA2 (double strand break repair), MSH6 and MGMT (both single strand break or point mutation repair). Differential equations describing the change of DNA damage over time were defined as follows:

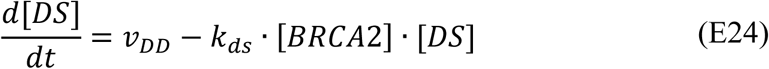

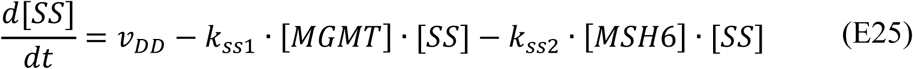

#### Initial conditions

The initial conditions of the DNA damage submodel were taken directly from their original source, except for the levels of repair enzymes, which came from proteomics measurements. Species in the original model possessed arbitrary units. We found the best approximation of these to an absolute unit of measurement to assume they represented units of μM, which we then converted to units of nM, scaling all rate constants accordingly.

#### “Digital” p53 dynamics

As mentioned in the original text, p53 dynamics are thought to be “digital” instead of “analog.” Namely, that increases in DNA damage are thought to increase the number of p53 pulses in a single cell without inducing a change in pulse amplitude or width (Lahav et al., 2004). To achieve this behavior, we needed to render the p53 response to DNA damage even more ultrasensitive. The original model already possessed a cooperative hill function for the activation of p53 by ATR and ATM. We added an additional cooperative hill function for the activation of ATM and ATR by DNA damage. The rate laws describing the activation of ATM and ATR are now as follows:

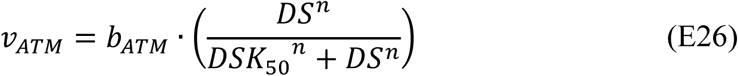

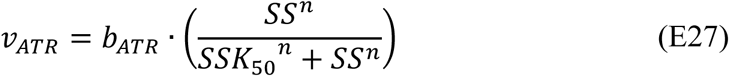

Here, *b*_*ATM*_ and *b*_*ATR*_ are rate constants and *DS* and *SS* are quantities of double-and single-stranded breaks in DNA. *DSK*_50_ and *SSK*_50_ (both set to 2nM; *n*=20) are EC_50_ concentrations for *DS* and *SS*, respectively (half-maximal concentrations required to induce a full response). This ensured a switch-like p53 response to DNA damage once a certain threshold of DNA damage was achieved. (In the original model, there was a DNA damage multiplier following the *b*_*ATM*_ and *b*_*ATR*_, with a value of either one or zero, serving as an on or off switch for DNA damage.) The rate of DNA damage repair also played an important role in ensuring that the number of p53 pulses increased with increasing DNA damage. As DNA damage is repaired and its levels drop beneath the threshold needed for activation of ATR and ATM, the activation of p53 is eventually ceased; however, while DNA damage remains above the threshold, highly regular p53 pulses persist (see Figure 3F and S3E in main text). Thus, in addition, we also manually fit several parameters of DNA repair (*k*_*ds*_, *k*_*ss*1_, and *k*_*ss*2_ from Equations E24 and E25) to enable this behavior.

### 2.5 Apoptosis submodel

*About*. The apoptosis submodel depicts the activation of initiator caspases (caspase 8 and 10) by TRAIL through the death receptors (DR4/5), which leads to the activation of executioner caspases (caspases 3 and 7), leading to cell death (Albeck et al., 2008). Initiator caspases can also activate a mitochondrial dependent pathway, driven by caspase 8 directed cleavage of Bid, which activates Bax. Active Bax can then oligomerize on the mitochondrial membrane and form pores leading to the release of cytochrome C. The release of cytochrome C from the mitochondria prompts the formation of the apoptosome, which activates caspase 9 (regulated by levels of XIAP and Smac). Caspase 9 can then feed back onto the activation of executioner caspases. There exists a positive feedback loop between initiator and executioner caspases (caspase 8 → caspase 3 → caspase 6 → caspase 8), which serves to create an all-or-nothing response once the appropriate levels of caspases have become activated (Eissing et al., 2004). Executioner caspases drive the digestion of many critical cellular components, which results in cell death. PARP, a DNA repair enzyme, is one target of caspase 3, and cleaved PARP (cPARP) is commonly readout for apoptosis (Tewari et al., 1995).

There are also intracellular modes of apoptosis signaling, through the regulation of pro- and anti-apoptotic proteins by phosphorylated ERK, phosphorylated AKT, PUMA, and NOXA. We added a few proteins to the original Albeck et al. (2008) model in order to describe intrinsic apoptosis signaling. We added BAD (pro-apoptotic Bcl2 protein, phosphorylated by ppERK and ppAKT) and BIM (pro-apoptotic Bcl2 protein, phosphorylated by ppERK and transcriptionally upregulated via ppAKT/FOXO). In addition, we added a bidirectional translocation reaction between cytoplasmic and mitochondrial forms of Bcl2 as it was not included in the original model.

#### Sensitivity and robustness of apoptosis submodel

The original apoptosis model created by Albeck et al. (2008) was designed to model extrinsic cell death processes in response to TRAIL over relatively short time scales. Because of this, they did not need to incorporate synthesis and degradation processes, or basal caspase activities. Because we are modeling over longer time scales, we added such mechanisms, which also allow robustness to small amounts of death signal. One mechanism by which the cell can do this is by rapidly degrading active, pro-apoptotic proteins (e.g. active caspases). We increased the degradation rates for several active pro-apoptotic proteins by several fold above their inactive form. Degradation of active TRAIL receptor complex was increased 1000 times and the degradation rates of active caspase 8, 3, 6, tBid, and active Bax were increased by 100 times above their inactive forms. Now, when a subthreshold amount of caspases becomes active, they are rapidly degraded, and the cell can return to homeostasis. Synthesis of all mRNAs and proteins in the model was incorporated via the expression submodel.

Once the MCF10A levels for apoptosis proteins were set, several rate constants needed to be modified to reduce the sensitivity of the model without losing the all-or-nothing response once a threshold of apoptosis signaling has been achieved. We made alterations to the reactions as described below in Table SI6. These changes are certainly not unique but do result in phenotypic predictions consistent with experimental observations.

**Table SI6.** Changes made to rate constants in original apoptosis model by Albeck et al (2008).

| Reaction | Fold change |
| --- | --- |
| Caspase 6 binding pro-caspase 8 | 1/1000 |
| Caspase 3 binding pro-caspase 6 | 1/100000 |
| Caspase 8 binding pro-caspase 3 | 1/100000 |
| Caspase 8 binding Bid | 1/10000 |
| Caspase 3 binding PARP | 1/10000 |
| Bax dimerization | 100000 |
| Bax2 (dimer) dimerization | 100000 |
| Bax2 (dimer) dissociation | 10000 |
| Bax4 (4-subunit protein) dissociation | 10000 |
| Apoptosome cleaving pro-caspase 3 | 1/10000 |
| Caspase 8 cleaving Bid | 1/10000 |
| koff for open pores | 1000 |
| Catalytic constant for open pores | 1/100 |
| Ligand-bound TRAIL receptor becoming active | 1/10 |

#### Intracellular apoptosis signaling

We desired that the model incorporate intracellular modes of apoptosis signaling, for which we required basal pro-apoptotic signaling. Intracellular modes of apoptosis are thought to rely primarily on the regulation of pro- and anti-apoptotic Bcl2 proteins. Anti-apoptotic Bcl2 proteins are thought to “keep the brakes” on cell death, releasing their inhibition of pro-apoptotic proteins only when an intracellular death signal is initiated or a pro-survival signal is abrogated. We reasoned that for this to be possible, a basal level of death signaling would have to be propagated through the pathway. This would lead to the creation of subthreshold levels of proapoptotic proteins (e.g., tBid), which would be sequestered by anti-apoptotic Bcl2 proteins until a death signal of sufficient magnitude and duration was received. To accomplish this, a basal amount of pro-caspase 8 cleavage was incorporated. This propagated a low amount of death signaling through the entire apoptosis pathway. The rate of basal caspase 8 cleavage was estimated as part of the initialization procedure (below), ensuring that it by itself did not cause apoptosis. In order to approximate the levels of cell death for serum-starved cells over a 72-hour time course, we increase the affinity of BIM for Bax by a factor (x5), post-initialization.

### 2.6 Cell cycle submodel

#### About

The cell cycle submodel is initiated by the upregulation of cyclin D mRNA by AP1 and cMyc. Cyclin D becomes active by binding to either Cdk4 or Cdk6 cyclin dependent kinases. Active cyclin D/Cdk4-6 phosphorylates Rb to de-repress E2F transcription. Subsequent upregulation of E2F causes the further upregulation of cyclin D and initiation of cyclin E and cyclin A induction (which bind to Cdk2 to become active), marking the beginning of S-phase and lasting into G2 phase. Finally, the activation of cyclin B/Cdk1 marks the beginning of mitosis, which, when complete, returns the cells to G1 phase.

Cell cycle equations are taken directly from their original source (Gérard and Goldbeter, 2009). We made a few changes to the original model. We removed the equations related to “ATR/Chk1 DNA replication checkpoint” except for the equations describing the upregulation of Chk1 in response to ATR (native to the DNA damage response submodel). We also removed the production of AP1 by growth factors, which is something that is now captured by the proliferation and growth submodel (through the formation of phosphorylated c-Fos bound to c-Jun).

#### Initial conditions

Because the original cell cycle model possessed intrinsic synthesis and degradation reactions, which we did not alter, we ran the model to a steady state in the absence of cycling and used these final (steady-state) species concentrations as the initial conditions. In addition, this submodel did not need mRNA quantities, as its synthesis reactions were already defined and we did not allow stochastic gene expression to affect it (see more below). We scaled all units from the original cell cycle model (species and rate constants) down by a factor of 10 to better agree with parameters from MCF10A cells.

#### Cell cycle entry driven by cyclin D mRNA

As described above, we removed the original growth factor terms and replaced them with the AP1 encoded by the Proliferation and Growth submodel. AP1 and MYC served as the primary proliferative inputs into the cell cycle model, whose levels cooperate to drive synthesis of cyclin D mRNA. The production of cyclin D above a certain threshold pushes the system beyond the “restriction point”, prompting the cascade of events initiating of the cell cycle and driving it to completion. Because flow cytometry data indicated that inputs from both ERK and AKT pathways were necessary for S-phase to proceed (Figure 6A), we modified the general form of the transcription equation (Equation E12) for cyclin D to reflect a need for both AP1 *and* cMyc (as opposed to AP1 *or* cMyc).

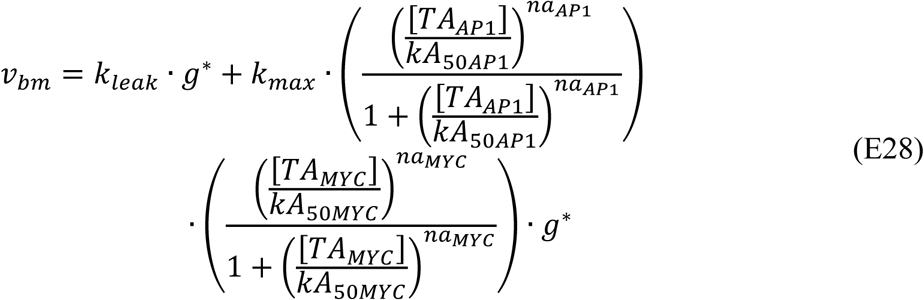

Here, *TA*_*AP*1_ and *TA*_*MYC*_ are the levels of transcriptional activators AP1 and MYC, respectively, and *kA*_50*AP*1_ (estimated to be 1.25 nM) and *kA*_50*MYC*_ (estimated to be 450 nM) are the half maximal concentrations of AP1 and MYC, respectively, and *na*_*AP*1_ and *na*_*MYC*_ (both set to *n*=3) are hill coefficients for AP1 and MYC induction, respectively. These parameters were manually estimated in order to match simulations to experimental data in terms of cyclin D induction (μ-Western blot data; Figure 6C) and percentage of cycling cells (BrdU flow cytometry data; Figure 6A) in response to EGF (10nM), Insulin (10μg/mL), and EGF + Insulin.

#### Order and timing of cyclin/cdk complexes

Importantly, this cell cycle model replicated the proper order and timing of cyclin/cdk complex oscillations (depicted in Figure SI2, below). As cyclin D becomes upregulated in G1 phase and binds to its cognate cdks, cdk 4 and 6, it provokes the phosphorylation of Rb, causing Rb to release its inhibition of E2F synthesis. An increase in E2F leads to the production of cyclin E and cyclin A, which define the entry into S- and G2-phase. These then lead to the activation of CDC25, the phosphatase responsible for activating cyclin B/cdk1, which defines entry into mitosis (M-phase). Post-mitosis, cyclin B/cdk1 levels return to baseline and mark re-entry into G1 phase or a possible exit into G0 phase if cyclin D levels have dropped considerably during cycling.

### Cell cycle duration

The simulated duration of one cell cycle event for cells treated with growth factors was approximately 20 hours, which is typical for a mammalian cell. This is also the approximate cell cycle duration measured for MCF10A cells (Albeck et al., 2013).

#### Inhibition of cycling by p21

Damage to DNA causes upregulation of p53, which is known to upregulate p21 (Riley et al., 2008), a potent cyclin dependent kinase inhibitor (CKI), which can bind to and sequester all cyclin/cdk complexes (Abbas and Dutta, 2009) leading to cell cycle arrest. In order to verify if our model could reproduce this qualitative behavior, we simulated DNA damage while cell cycle oscillations were present (cell cultured in growth factors) and looked for cell cycle arrest. As is clear from Figure SI3, below, upregulation of p21 results in the ceasing of cyclin oscillations and interrupts cell cycle progression. Also, when DNA damage is repaired (by DNA repair enzymes), and p53 activation returns to baseline, the cell is able to re-enter the cell cycle.

#### Unresolved emergent fragility of cell cycle to expression noise

As described above, every gene product in our model is linked to stochastic gene switching and mRNA production/degradation, giving rise to temporal fluctuations in protein levels. When we attempted to link the cell cycle submodel to stochastic gene expression, we obtained non-sensical simulation results, including rampant spontaneous cycling in the absence of Cyclin D induction, spurious expression of the various cyclins with incorrect ordering, and a lack of regular frequency, amplitude and duration of cyclin peaks. This fragility of cell cycle models to stochastic gene expression-like noise is documented by previous studies (Kar et al., 2009). This observation implies there are as-yet undiscovered cellular mechanisms that provide natural robustness of the cell cycle to gene expression (or other) noise, solving which is outside the scope of this manuscript. Therefore, we retained a deterministic formulation of the cell cycle sub-model.

### 2.7 Submodel integration

Links between submodels were created based on well-studied mechanistic interactions that are known to exist between proteins in each submodel (see Figure S1 for a detailed mechanistic schematic).

#### Survival Signaling—Proliferation and Growth to Apoptosis

The primary species that mediate survival signals in the model are active ERK and Akt kinases (ppERK and ppAkt). ERK is known to phosphorylate the pro-apoptotic protein BIM, inactivating its ability to active Bax (Hübner et al., 2008). AKT is known to phosphorylate FOXO (Zhang et al., 2011), which when phosphorylated translocates into the nucleus to induce transcription of target genes. These target genes include receptor tyrosine kinases (see below) and BIM (Gilley et al., 2003), which can active Bax. Both ERK and AKT are known to phosphorylate BAD (Fang et al., 1999; del Peso et al., 1997), which renders it inactive and unable to bind to, and sequester, anti-apoptotic Bcl2 proteins.

#### G0/G1 Transition—Proliferation and Growth to Cell Cycle

Signal flux through the ERK and AKT pathways can also induce the cell cycle by upregulating cyclin D. Specifically, cyclin D is upregulated by AP1 (Shaulian and Karin, 2001) (phosphorylated cFos in complex with cJun) and cMyc (Bouchard et al., 1999), induced by the ERK and AKT pathways, respectively. Moreover, AP1 induces transcription of cJun, leading to positive feedback, which further amplifies AP1’s influence on cyclin D (Angel et al., 1988).

#### Intrinsic Death—DNA Damage to Apoptosis

In a non-transformed cell, the induction of DNA damage promotes apoptosis if the damage is not repaired in a timely fashion (Roos and Kaina, 2006). The cell does this to protect against the potential for developing disease, such as cancer, as a result of the DNA damage. Here, we allow active p53 to upregulate PUMA (Nakano and Vousden, 2001) and NOXA (Oda et al., 2000), both proapoptotic proteins that bind to, and sequester, anti-apoptotic Bcl2 proteins.

#### Cell Cycle Arrest—DNA Damage to Cell Cycle

When a non-transformed cell sustains DNA damage, it arrests the cell cycle in order to prevent the spread of potentially deleterious mutations to daughter cells. This is thought to occur primarily through the transcriptional upregulation of p21 by active p53 (Riley et al., 2008). p21 is a cyclin dependent kinase inhibitor which binds to and inactivates all cyclin/cdk complexes (Abbas and Dutta, 2009). The other mechanism that we encode in our model is the upregulation of Chk1 by active ATR (Hekmat-Nejad et al., 2000), which is activated primarily in response to single-stranded breaks. Chk1 promotes cell cycle arrest by phosphorylating CDC25 proteins, which targets it for rapid degradation. Chk1 can inhibit all CDC25 isoforms (Karlsson-Rosenthal and Millar, 2006). CDC25 are phosphatases that activate many cyclin/cdk complexes. For example, it removes an inhibitory phosphate group from cyclin B/cdk1, rendering it fully active which promotes the initiation of mitosis.

#### Feedback on Survival Signaling—Proliferation and Growth to RTK

As is commonly seen across numerous signaling pathways, activation is usually followed by a phase of deactivation or negative feedback, preventing chronic and potentially deleterious pathway activation. Signaling through the ERK pathway is known to affect receptor tyrosine kinases by phosphorylating them on specific sites that blunt their ability to signal downstream (Li et al., 2008), providing a mechanism of negative feedback. In our model ERK phosphorylates and inactivates the ErbB family members (ErbB1-4). There are mechanisms of transcriptional feedback as well. Receptor tyrosine kinases are known transcriptional targets of FOXO (we allow FOXO to transcriptionally upregulate all receptor tyrosine kinases in our model), which translocates into the nucleus upon dephosphorylation; when AKT is active, FOXO is phosphorylated and cytoplasmic (Tzivion et al., 2011; Zhang et al., 2011). Sprouty proteins are transcriptionally upregulated by active ERK, which bind to and inactivate multiple RTKs (Ozaki et al., 2001).

#### Protein synthesis—Expression to Other Submodels

Synthesis processes are captured by the expression submodel, which controls the production of proteins across most of the submodels (with the exception of the cell cycle, as described above). We model 141 total genes. These genes are translated to create only 102 “protein conglomerates”, as some of the gene isoforms perform redundant functions or are experimentally indistinguishable (see Supplementary Table 1 for mapping between proteins and gene isoforms). Functionally redundant isoforms are summed into the final protein amount, *p*, which we term a conglomerate:

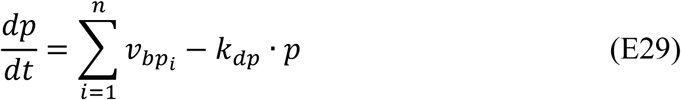

Here, 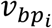 is the translation rate for every functionally redundant isoform, *i*, for protein, *p*. For example, the ERK protein is functionally composed of two main isoforms: MAPK1 (ERK2) and MAPK3 (ERK1) (Figure 2B). Upon translation, these two isoforms are summed to create the protein conglomerate “ERK” in the model.

## Chapter 3: Initialization Procedure

### 3.1 Initialization rationale, approach, and output

The goal of the initialization procedure is to ensure that the steady state levels for all proteins and mRNAs in the model match what we have measured in serum-starved MC10A cells. It is also done to capture phenotypic behavior that one would expect for cells in the serum-starved state (e.g. not cycling, dying in small numbers, etc.). The model commences with protein quantities given to species that represent proteins that have been immediately translated from mRNA; that is, they lack any post-translational modifications. However, the levels of post-translationally modified proteins may change when we run the model due to basal activities (e.g. basal Ras, PIP_3_, and caspase 8 activity; more below) or other phenomena (e.g. phosphorylation of β-catenin by GSK3-β; formation of heterodimeric complexes). These post-translationally modified forms of proteins may have different degradation rates than their nascent counterparts. Thus, if run to steady state, the total amount of a given protein may no longer equal what was measured via proteomics.

To correct for this, we increase or decrease synthesis rates by altering gene-specific translation rate constant (*k*_*TL*_) values. We do this using an iterative approach. Every time we introduce a new element to the system (e.g. such as different basal activities), we run the model to an effective steady state in the absence of growth stimulus (1000 hours) and calculate a ratio between simulated (sum across all species that possess a given protein) and measured protein levels. We then multiply the translation rate constant (*k*_*TL*_) by this ratio and re-run the simulation. We repeat this process until simulations are within 1% of experimental data. All simulations during initialization are performed deterministically.

The process of initializing transcription is described below under subheading TARs, however the goal is the same (post-initialization mRNA levels should match mRNA-seq measurements). Quantification of the final initialization results can be found in main text Figure 2E. Overview of initialization process can be found in Figure SI4, below.

### 3.2 Step 1: Basal Ras and PIP activities

The first step of initialization begins with all post-translationally modified forms of proteins set to zero. During this step, basal activity fluxes are turned on. Basal fluxes through the ERK and AKT pathways are encoded by two activities: (*i*) basal Ras activation and inactivation reactions (e.g. intrinsic RasGTPase activity) and (*ii*) basal PIP2 phosphorylation and dephosphorylation reactions. Here, we employ the iterative approach described above to alter the translation rate constants (*k*_*TL*_) until steady-state protein levels match experimentally-measured protein levels. Importantly, during this step we prohibit EIF4E from affecting the overall translation rate. We do this because basal fluxes as well as the binding of EIF4E to mRNAs induce large changes in free EIF4E levels. Because EIF4E can alter translation rates globally, rapid changes in its levels make it difficult to approximate a steady state for each species. We accomplish this by keeping the free EIF4E levels in the translation equation from Equation E12 set to total EIF4E levels (as measured by proteomics), so that changes in free EIF4E cannot affect the overall translation rate. This converts the translation equation to:

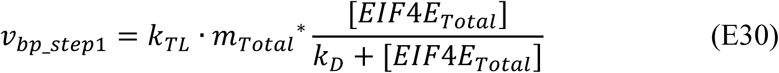

Once good agreement between simulation and experiment is achieved, a final free EIF4E quantity is defined. Using this quantity, we calculate a new *v*_*bp*_ for each gene and multiply the translation rate constants (*k*_*TL*_) by the ratio between the *v*_*bp*_*step*1_ and *v*_*bp*_.

### 3.3 Step 2: Basal cyclin D synthesis and p21 degradation

This initialization step is performed in order to estimate two parameters in the cell cycle submodel. As mentioned before, synthesis and degradation reactions for the cell cycle submodel are intrinsic to that submodel. We also keep the initial concentrations from the original model, with the exception of cyclin D and p21, which we set using our proteomics data. To set initial cyclin D levels we estimate a basal cyclin D synthesis constant. The rate equation describing cyclin D synthesis is as follows:

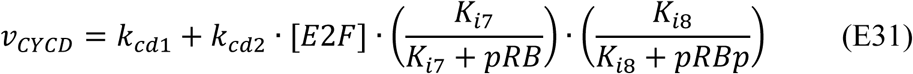

The first term, *k*_*cd*1_, is a zero-order synthesis for cyclin D driven by basal levels of transcriptional activators (Gérard and Goldbeter, 2009). The second term describes cyclin D synthesis as driven by E2F levels and inhibited by Rb levels. In this initialization step, we estimate the value of *k*_*cd*1_. We employ a similar iterative approach as described above until the total amount of cyclin D in the system is equal to the amount measured via proteomics. Importantly, we ensure that this amount of cyclin D does not induce cell cycle oscillations, as the cells do not cycle in the serum-starved state. To set p21 levels we estimate the degradation rate constant for p21, which follows first-order degradation kinetics. We do this in the same iterative manner as for cyclin D.

### 3.4 Step 3: Basal caspase 8 cleavage

The ability of an internal pro-apoptotic stimulus (e.g. PUMA/NOXA upregulation by p53) to induce apoptosis depends in large part on the negative regulation of anti-apoptotic proteins. This is only possible if there is a basal flux of death signaling through apoptotic pathways. Indeed, if there was no basal death signaling, there would be no pro-apoptotic proteins for the anti-apoptotic Bcl2 proteins to bind, and the inhibition of anti-apoptotic Bcl2 would do nothing to induce cell death. Basal death signaling could be the result of basal caspase activity, which is known to occur in the absence of an overt death stimulus (Gdynia et al., 2007; Marini et al., 2014). We chose to add a basal, first-order caspase 8 cleavage reaction to simulate this effect. Prior to this initialization step, the basal caspase 8 cleavage rate is set to zero. In this initialization step, we iterate through increasing values (log-spaced) of the caspase 8 cleavage rate constant until apoptosis occurs within a 1000-hour simulation time window. We choose the value that is one increment lower than the value that caused apoptosis. Finally, because the introduction of this rate may render a mismatch between the quantities of various proteins in the model and their experimentally measured values, we also re-approximate translation rate constants (as above) until all model protein levels match their experimentally measured values. We then test to make sure the cell still undergoes apoptosis as previously.

### 3.5 Step 4: Basal DNA damage

Due to insults from the environment as well as internal stressors, there exists a low, basal amount of DNA damage inside living cells even when no explicit insult is present. The effects of basal DNA damage are enhanced during S-phase, as the DNA is uncoiled from histones and becomes more venerable to insult. In addition, because DNA is being shuffled so much during replication, breaks are more likely to occur. However, this basal DNA damage plus replicative stress should not be enough to induce a p53 response on average (i.e. deterministic regime). The goal of this initialization step is to calculate a level of basal DNA damage activity (described by the term *k*_*basal*_, in Equation E23) such that (*i*) the total amount of active p53 is between 1-5% of its total quantity and that (*ii*) a full p53 response is not elicited by the basal activity. To do this, we estimate a value for *k*_*basal*_ such that 1-5% of p53 is active and such that the quantity of single- and double-stranded DNA breaks does not exceed its half-maximal effective concentration (*SSK*_50_ and *DSK*_50_, respectively, from Eqs E5 and E5, below) needed to induce full activation of ATR and ATM, respectively. The rate law for the production of DNA damage (*v*_*DD*_) is defined above in Equation E23, and placed here for convenience:

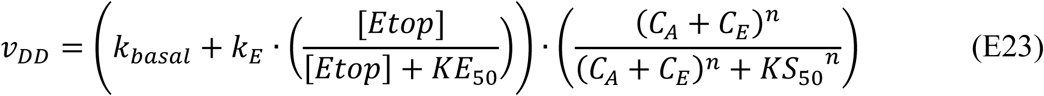

Recall the differential equations for double- and single-stranded breaks from above:

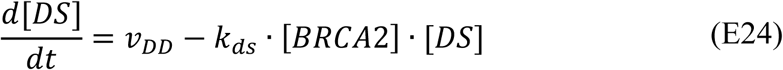

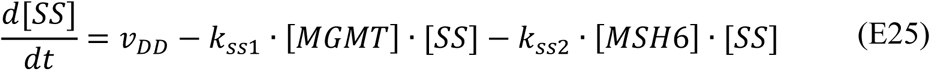

When a cell is in S-phase and untreated with Etoposide, *v*_*DD*_ is equivalent to *k*_*basal*_ (basal DNA damage rate). Solving Equations E24 and E25 above at steady state for *DS* and *SS* gives:

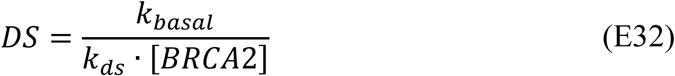

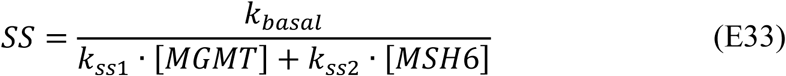

Thus we ensure that the quantity for *DS* and *SS* do not exceed *DSK*_50_ and *SSK*_50_ (both set to 2nM) in the following rate laws describing activation of ATM and ATR from above, repasted here for convenience:

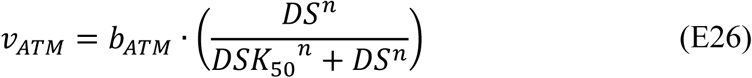

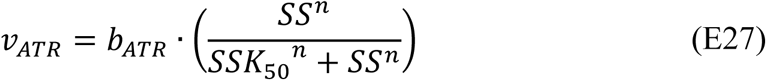

Our final active p53 percentage was approximately 2% and this did not induce a full p53 response.

### 3.6 Step 5: Transcriptional activators and repressors (TARs)

The goal of this initialization step is to ensure that mRNA synthesis rates are in equilibrium with mRNA degradation rates, particularly for mRNAs that have modeled regulation. These transcriptional activators and repressors (TARs) that perform such regulation are the species that transfer information from the deterministic model to the stochastic model. As depicted in Equation E12, above, the differential equation for mRNA production contains the rate of transcription, which possesses a “leak” or constitutively active term plus an “induced” or TAR induced expression term, minus the rate of mRNA degradation. The general form of the differential equation is as follows:

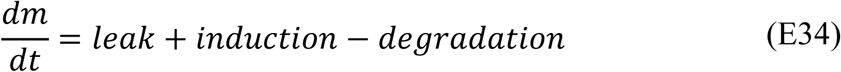

The rate of mRNA degradation was already known, calculated from mRNA half-lives and mRNA levels as described above. We then simply required that, at steady-state, the rate of transcription was equal to the rate of degradation:

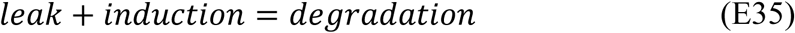

We first calculated the induction term, which, as depicted in Equation E12, is a function of the levels of TARs, and preset K_50_ values and hill coefficients. The level of each TAR is taken from the fully equilibrated model. K_50_ values and hill coefficients are estimated to (*i*) ensure that induction plus leak terms do not exceed degradation terms and (*ii*) to fit the model to experimental results or other empirical observations. For example, the parameters for induction of cyclin D by AP1 and cMyc were estimated to approximate cyclin D protein induction as measured by μ-Western blot and percentage of cycling cells in response to different growth factors as measured by BrdU incorporation flow cytometry experiments. Leak terms are then calculated to balance out the equation; specifically the *k*_*leak*_ term from Equation E5. If a gene did not possess any explicitly coded TARs, the induction term was zero, and the leak term was equal to the degradation term.

## 4. Supplementary Figure Legends

**Figure SI1.**
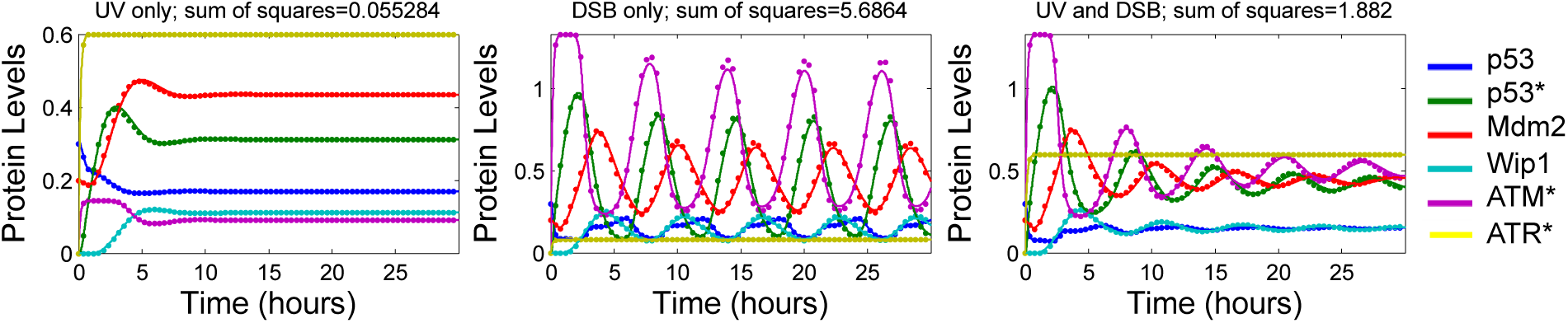
Simulation results comparing original delay differential equation model (Batchelor et al., 2011) displayed with dots and ordinary differential equation model fit displayed with lines in same color.

**Figure SI2.**
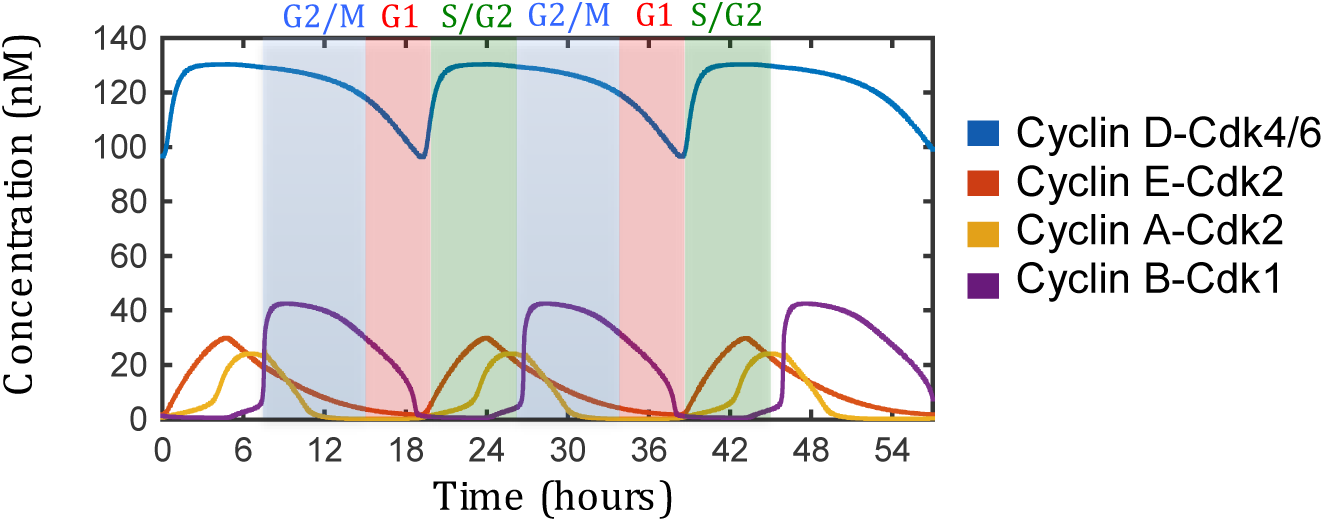
Timing and order to cyclin/cdk complexes and their correspondence with cell cycle stage.

**Figure SI3.**
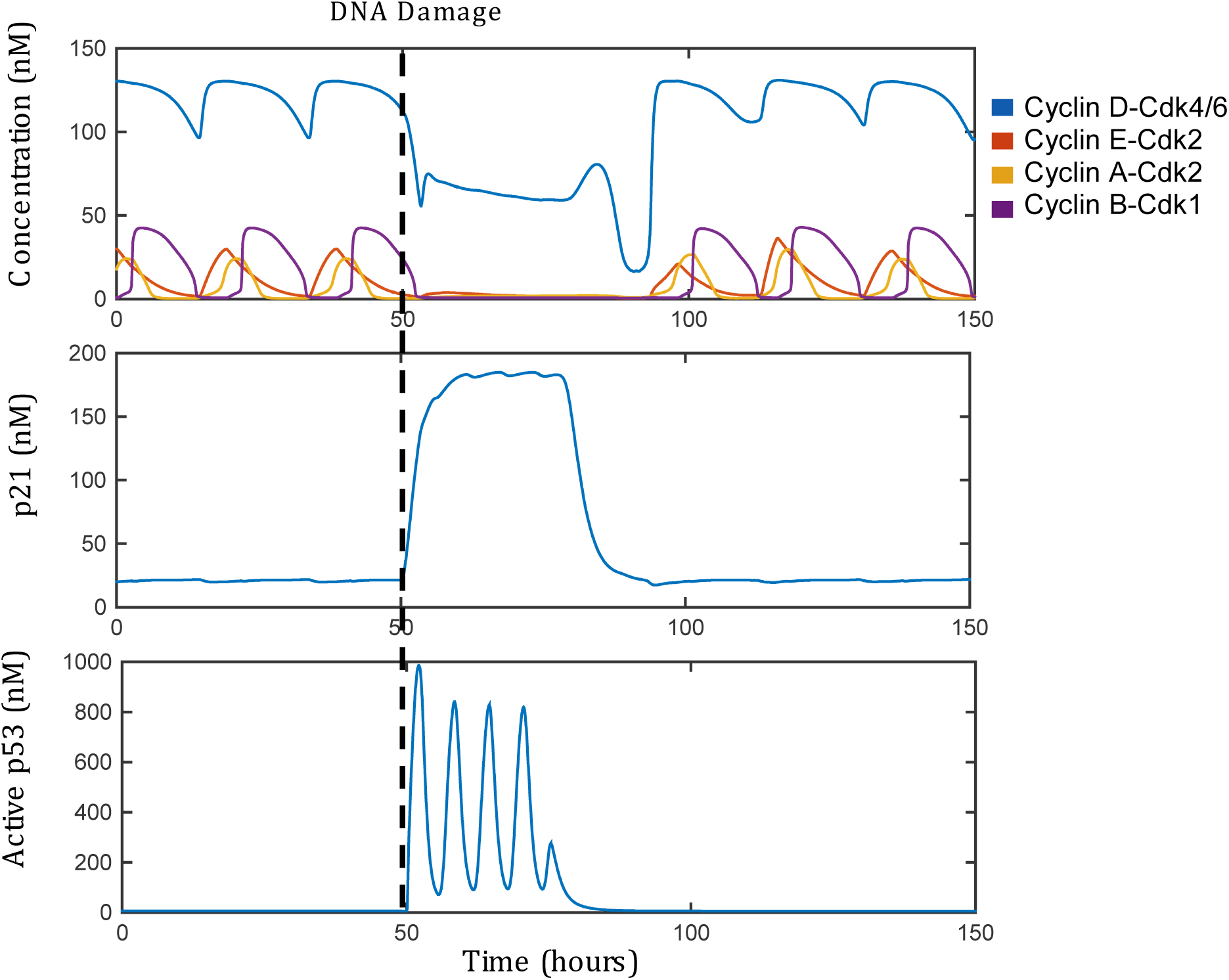
DNA damage is induced at 50 hours for a cycling cell. The activation of p53 (bottom) causes upregulation of p21 (middle), which arrests the oscillatory behavior of cyclic/cdk complexes (top).

**Figure SI4.**
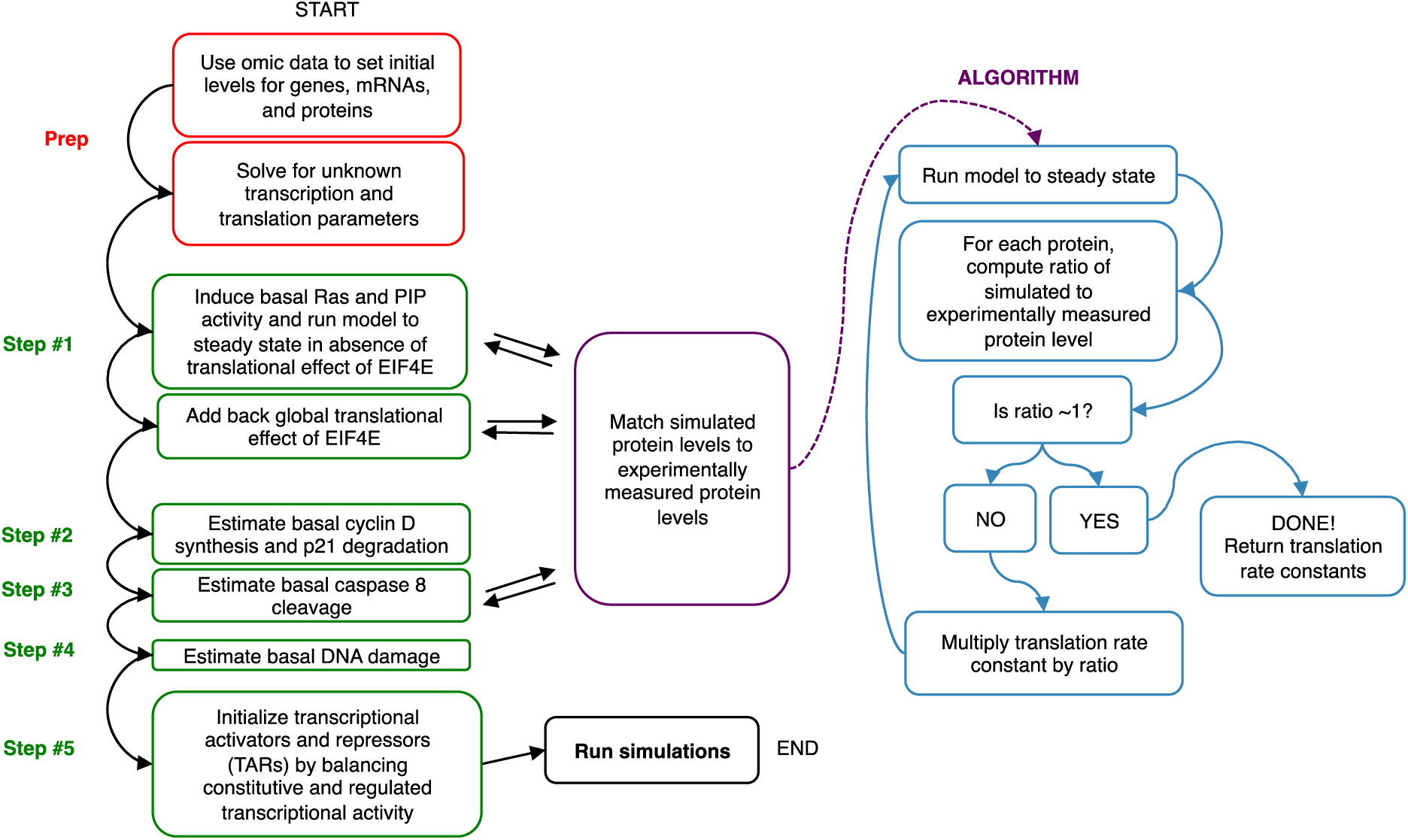
Schematic depicting overview of initialization process.

**Figure S1.**
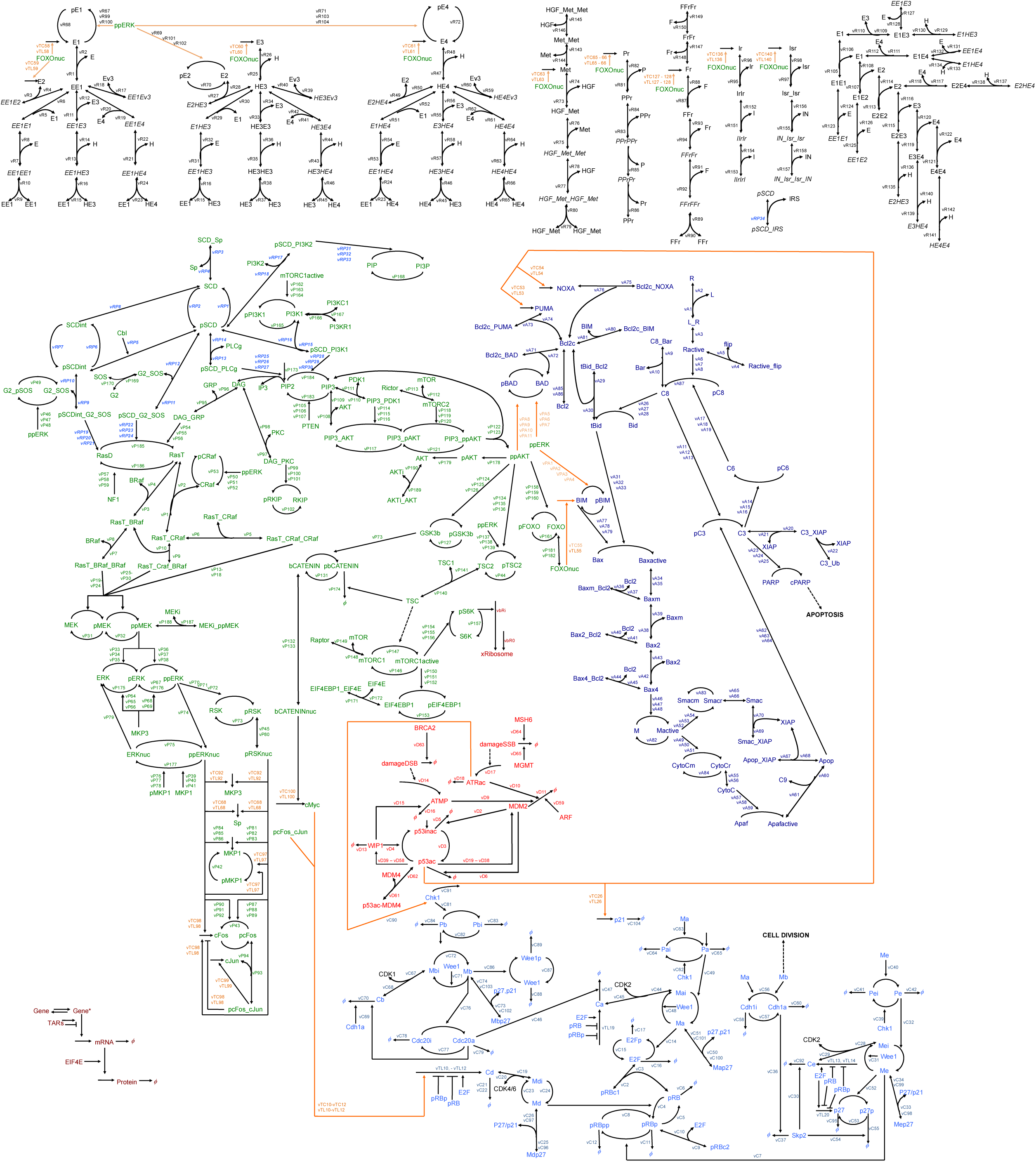
Detailed kinetic scheme of model. This kinetic scheme depicts all of the reactions in the model. Labels correspond to the indices for each rate law in the equations and code. A “p” prior to a protein name indicates it is phosphorylated. An underscore between two proteins indicates they are in complex. An arrow pointing to a curved arrow indicates three reactions: association, dissociation, and catalysis. A dashed arrow indicates a lack overt mechanism, described with either Hill or Michaelis-Menten rate laws.

**Figure S2.**
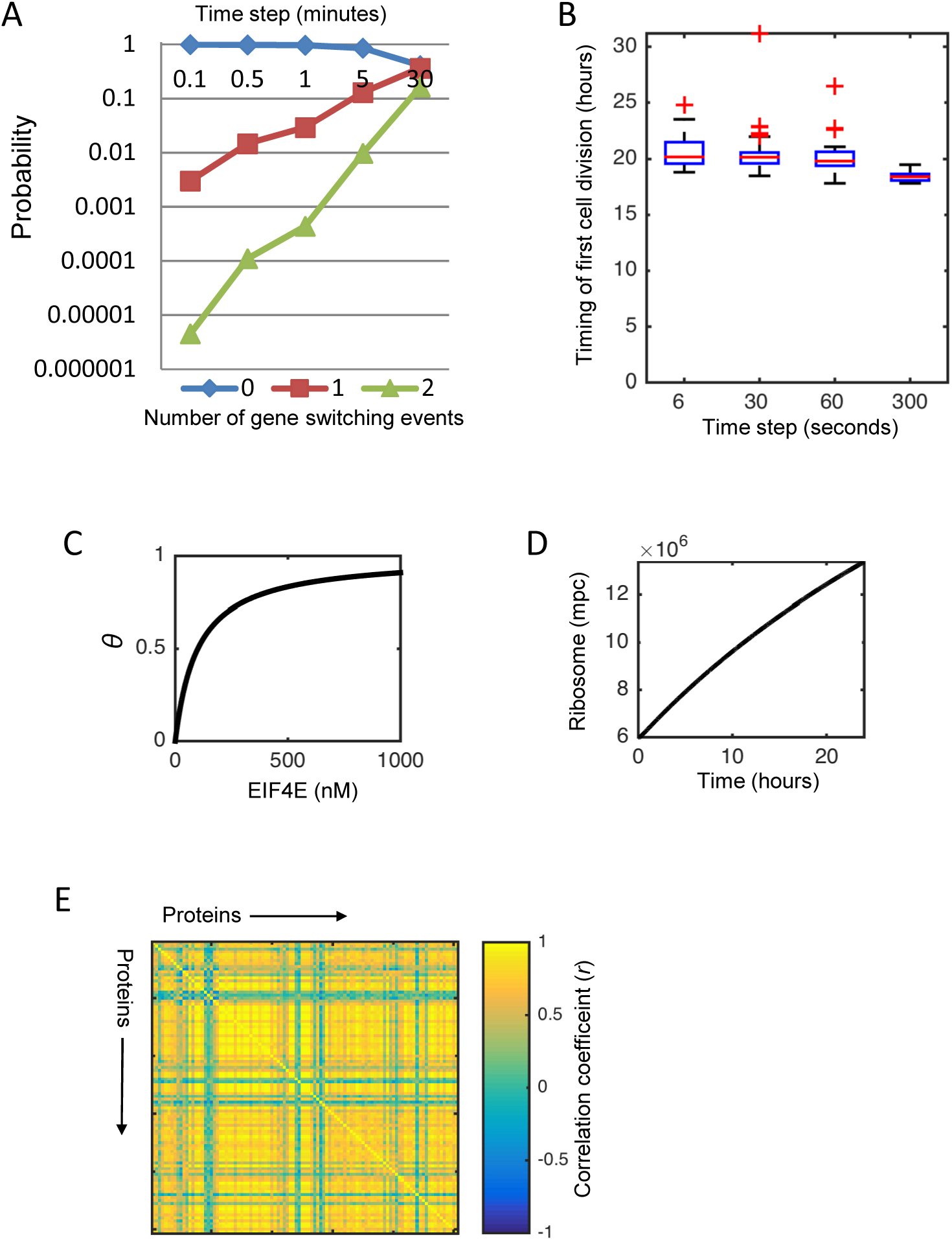
Properties of the Expression Submodel. A) Probability of zero (blue diamonds), one (red squares), or two (green triangles) gene switching events given different time step intervals between the stochastic and deterministic components of the model. We chose 30 seconds as a tradeoff. B) Cell-to-cell variability in the timing of the first cell division event following growth factor stimulation. Different time steps between the stochastic and deterministic components of the model are shown along the x-axis. Growth factor stimulation is with all growth factors set to 10nM, except for insulin, which was set to 1721nM. Variability seems rather consistent for the 6, 30, and 60 second time steps, though it collapses when the time step reaches 300 seconds. C) The solution to the hill function describing the effect of EIF4E on translation rate. D) The number of ribosomes in a single cell doubles during the course of a typical cell cycle duration (∼20 hours) in response a full mitogenic stimulus (same as B). E) Correlation between protein levels across a population of cells, showing the typically observed global positive correlations.

**Figure S3.**
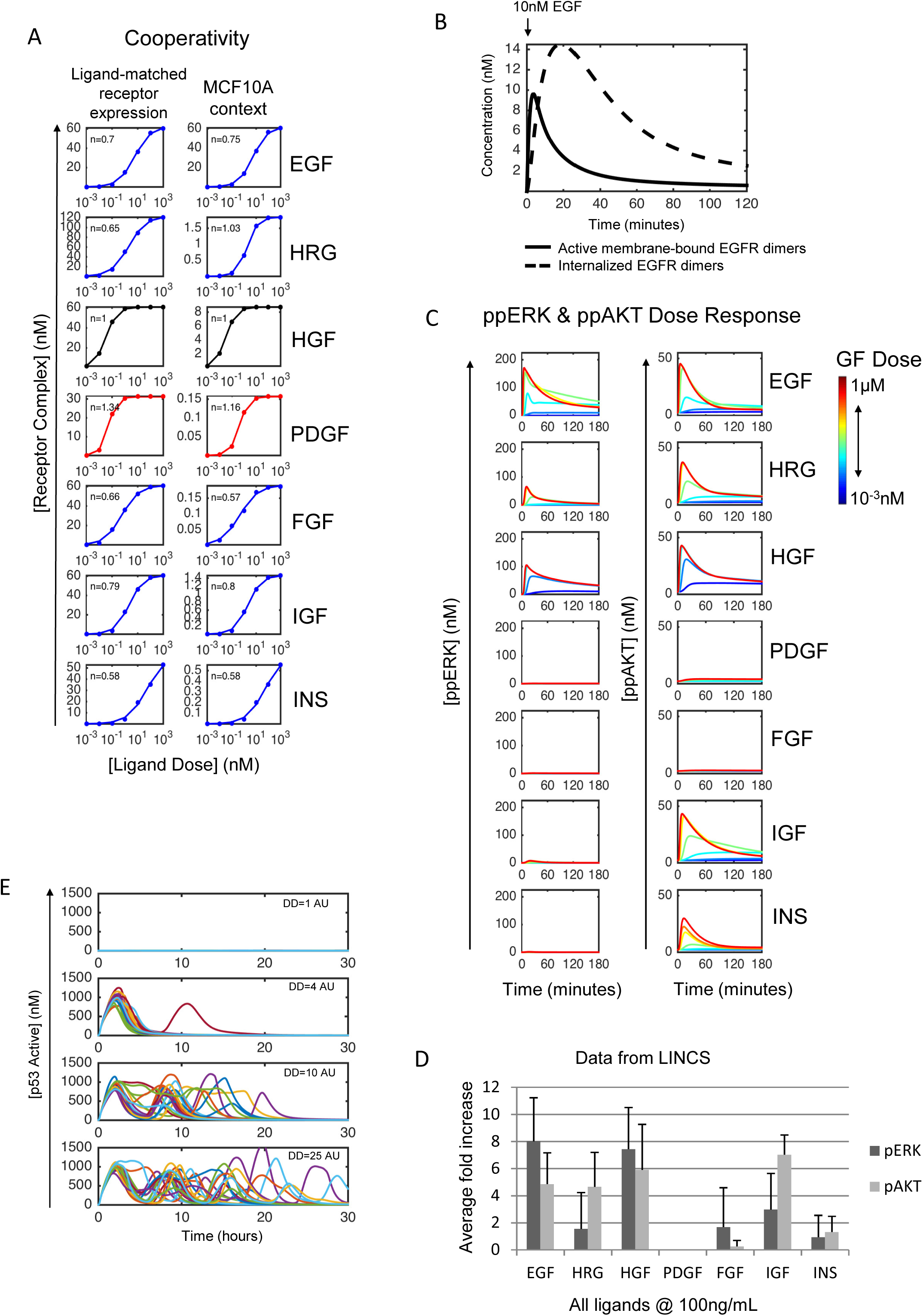
Additional Unit Testing. A) Cooperativity profiles comparing the ideal case (left)—when only the receptors that bind each ligand are present on the cell surface and at appreciable levels—to those for the model tailored to expression levels for MCF10A cells (right). B) Time course plot showing dynamics of active plasma membrane-bound EGFR dimers (solid black line) and internalized receptors (dashed black line) in response to 10nM EGF, showing kinetics largely consistent with prior knowledge on trafficking and downregulation. C) Dose response curves for ppERK and ppAKT in the MCF10A-tailored model in response to various ligands at doses ranging from 1μM to 10^−3^nM. D) Data obtained from the LINCS (Library of Integrated Network Cellular Signatures) project (Niepel et al., 2014) showing average (average across 10, 30, and 90 minute time points) levels of phosphorylated ERK (dark grey) and AKT (light grey) in response to ligands depicted in model. Relative activation amplitudes are comparable to model predictions from Figure S3C, although IGF induces more ERK activation than predicted by our model. E) Dynamic data from the stochastic simulations that comprise Figure 3F (different cells are different colors). DNA damage increases from top to bottom. Although the number of pulses increases with increasing DNA damage, pulse height and width remain relatively constant.

**Figure S4.**
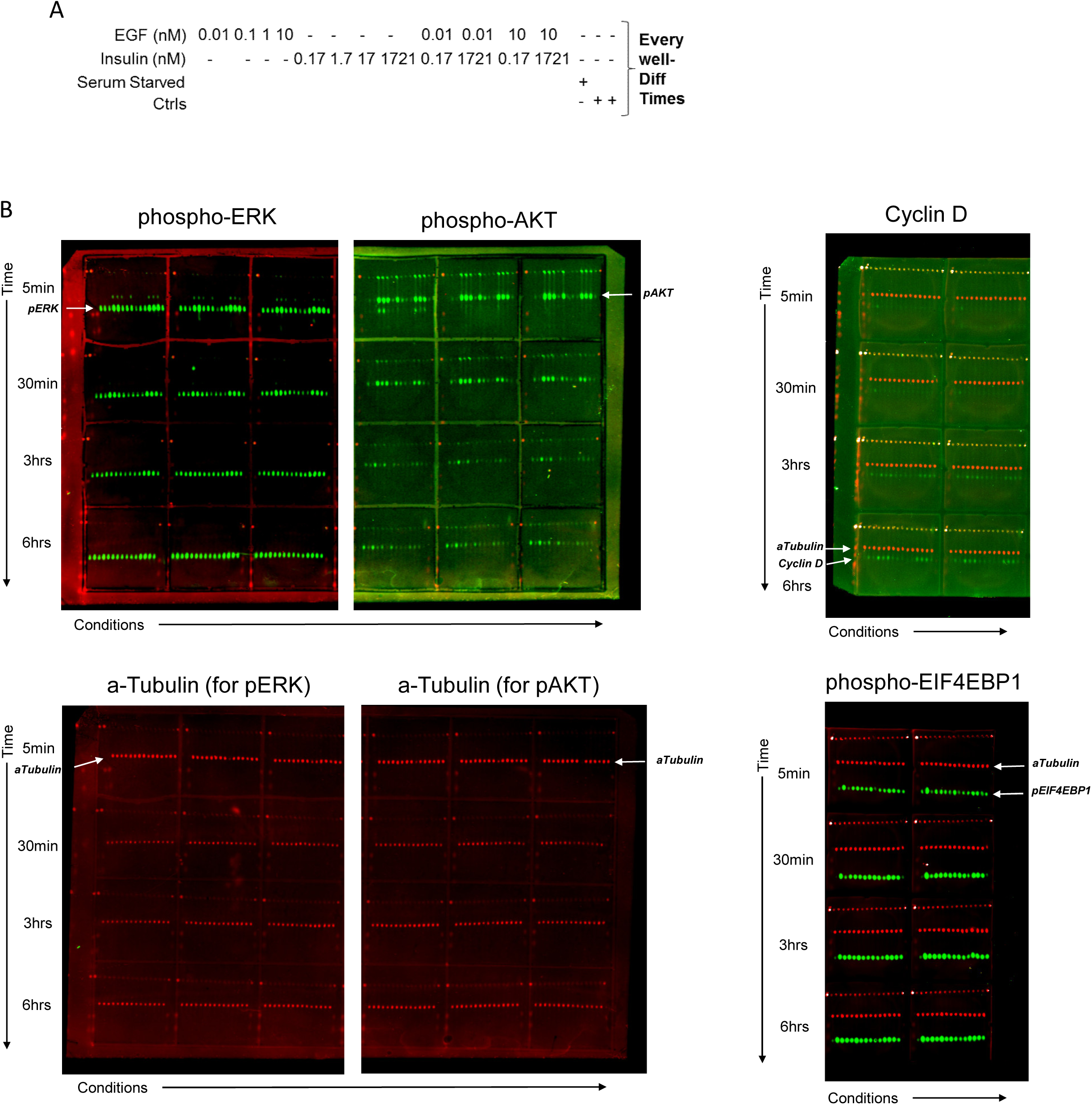
μ-Western blot raw images. A) Schematic outlining layout of conditions on μ-Western blot for every well. Concentrations are in units of nM. B) Raw scans of μ-Western blot membranes probed for pERK, pAKT, pEIF4E-BP1, α-Tubulin, and cyclin D (see Methods). Time points go from top-to-bottom. Treatment conditions go from left-to-right, as indicated in (A). Each row has biological replicates (three for pERK and pAkt, two for Cyclin D and p4EBP1).

**Figure S5.**
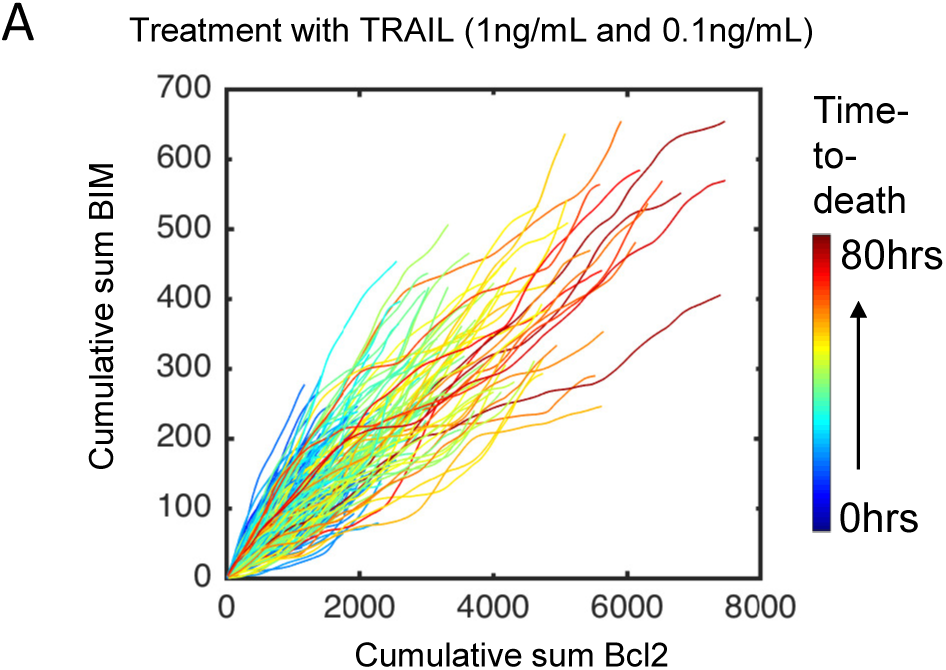
Predicting Apoptosis for Extrinsic Death. A) Simulated cells are treated with low doses of TRAIL (1ng/mL and 0.1ng/mL) and plots show the cumulative sum of BIM and Bcl2 levels across the time course during which they are alive (analogous to Figure 5F). Cell trajectories are colored in terms of their time to death. Here we see no obvious relationship between BIM and Bcl2 that is dependent on time to death as we saw in Figure 5F, highlighting to the stimulus-specific nature of apoptosis sensitivity.

**Figure S6.**
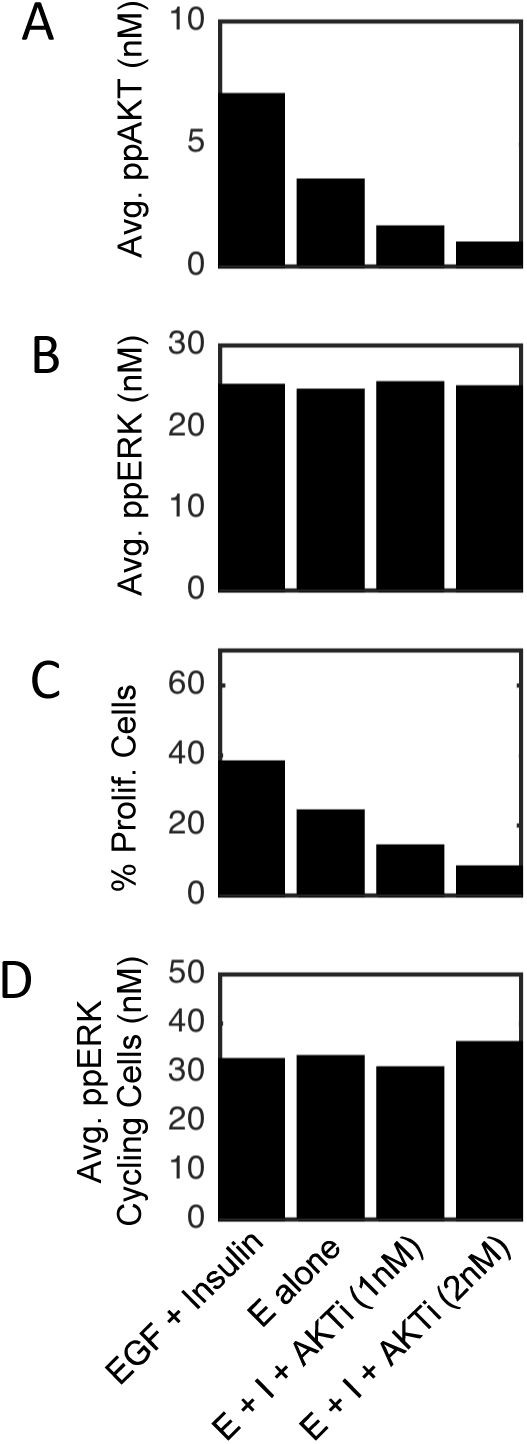
Time-averaged ERK and AKT activities in simulations. A) The average, time-integrated ppAKT levels across different types of stimuli—EGF (10nM) + Insulin (10μg/mL), EGF (10nM) alone, EGF (10nM) + Insulin (10μg/mL) + small AKTi dose (1nM), or EGF (10nM) + Insulin (10μg/mL) + small AKTi dose (2nM). A decrease in ppAKT is observed. B) The average, time-integrated ppERK levels across the same stimuli as in (A). No decrease in ppERK is observed. C) The percentage of proliferating cells at 24 hours post stimulation reveals that diminishing amounts of ppAKT signal decreases the percentage of proliferating cells. D) The average, time-integrated ppERK levels for cycling cells reveals no relationship between ppERK levels in cycling cells and ppAKT levels.

